# Pore constrictions in intervessel pit membranes reduce the risk of embolism spreading in angiosperm xylem

**DOI:** 10.1101/2020.10.19.345413

**Authors:** Lucian Kaack, Matthias Weber, Emilie Isasa, Zohreh Karimi, Shan Li, Luciano Pereira, Christophe L. Trabi, Ya Zhang, H. Jochen Schenk, Bernhard Schuldt, Volker Schmidt, Steven Jansen

## Abstract

- Embolism spreading in angiosperm xylem occurs via mesoporous pit membranes between vessels. Here, we investigate how the size of pore constrictions in pit membranes is related to pit membrane thickness and embolism resistance.
- In three models, pit membranes are modelled as multiple layers to investigate how pit membrane thickness and the number of intervessel pits per vessel determine pore constriction sizes, the probability of encountering large pores, and air-seeding. These estimations were complemented by measurements of pit membrane thickness, embolism resistance, and number of intervessel pits per vessel (*n* = 31, 31, and 20 species, respectively).
- Constriction sizes in pores decreased with increasing pit membrane thickness, which agreed with the measured relationship between pit membrane thickness and embolism resistance. The number of pits per vessel affected constriction size and embolism resistance much less than pit membrane thickness. A strong relationship between estimated air-seeding pressures and measured embolism resistance was observed.
- Pore constrictions provide a mechanistic explanation why pit membrane thickness determines embolism resistance, and suggest that hydraulic safety can be uncoupled from hydraulic efficiency. Although embolism spreading remains puzzling and encompasses more than pore constriction sizes, angiosperms are unlikely to have leaky pit membranes, which enables tensile transport of water.

## Introduction

Xylem sap in vessel-bearing angiosperms crosses numerous intervessel walls from the root to the leaf xylem, depending on the plant size, vessel length, and intervessel connectivity. An average angiosperm vessel is estimated to have about 34,000 intervessel pits, with values for different species varying more than 200-fold, from ca. 500 pits to > 100,000 (sample size, *n* = 72 species; Fig. S1). Each bordered pit pair has a pit membrane, which is mainly composed of ca. 20 nm wide cellulose microfibril aggregates. These pit membranes develop from the primary cell wall and middle lamella, and have a mean diameter of 4.8 ± 2.4 μm (*n* = 43 species; Jansen et al., 2009, 2011). Before pit membranes become hydraulically functional, hemicellulose and pectin compounds are enzymatically removed (O’Brien, 1970; Herbette *et al.*, 2015; Klepsch *et al.*, 2016). Therefore, fully mature pit membranes are non-woven, fibrous porous media of mainly cellulose, with a thickness between ca. 160 and 1,000 nm (Esau 1977; Pesacreta *et al.*, 2005; Kaack *et al.*, 2019).

Intervessel pit membranes play an important role in plant water transport by providing ca. 50% of the total hydraulic xylem resistance (Choat *et al.*, 2008). They control the immediate entry of gas from neighbouring, embolised conduits, and may become sites of further embolism propagation under persistent drought (Zimmermann, 1983; Brodersen *et al.*, 2013; Choat *et al.*, 2016; Brodribb *et al.*, 2016; Roth-Nebelsick, 2019). The lack of a mechanistic understanding of gas bubble movement through pit membranes, which is described as “air-seeding”, represents one of the major knowledge gaps in our understanding of water transport in plants (Jansen *et al.*, 2018). It is known that propagation of drought-induced embolism from one vessel to a neighbouring vessel is affected by pore dimensions of intervessel pit membranes, but how pit membrane thickness (*T*_PM_; see Table 2 for an overview of the acronyms used) affects pore dimensions and embolism spreading is unclear.

Instead of perfectly flat, two-dimensional structures, as often portrayed in textbooks, pit membranes are porous media with pores that include multiple constrictions, with the respective narrowest constriction in each pore governing flow of water and gas (Fig. 1; Kaack *et al.*, 2019) and, consequently, embolism spreading. Estimates of bottleneck diameters, *i.e.* constriction sizes, vary from 5 nm to well above 200 nm (Fig. 1; Choat *et al.*, 2003; Sano, 2005; Jansen *et al.*, 2009; Hillabrand *et al.*, 2016). Part of this variation is caused by sample preparation for imaging by scanning electron microscopy (SEM), which induces up to 50% shrinkage of *T*_PM_ during drying, with frequently enlarged pores and cracks (Shane *et al.*, 2000; Jansen *et al.*, 2008; Zhang *et al.*, 2017). Moreover, the challenge is to quantify size and shape of pit membrane pores in a three-dimensional approach. A three-dimensional model based on transmission electron microscopy (TEM) of fresh and shrunken pit membranes indicated a high porosity (*i.e.* void volume fraction) of 81%, highly interconnected pore systems with non-tortuous, unbending passageways, a lack of dead-end pores, and the occurrence of multiple pore constrictions within a single pore (Zhang *et al.*, 2020). Based on a shrinkage model and gold perfusion experiments, we found that constriction sizes in pit membrane pores vary from 5 to < 50 nm, with an average diameter around 20 nm (Choat *et al.*, 2003, 2004; Zhang *et al.*, 2020). Moreover, the structural characteristics of pit membranes appear to be fairly constant for angiosperm species, despite considerable variation in *T*_PM_. Indeed, pore constriction sizes around 20 nm occur both in species with thin (ca. 200 nm) and thick (> 500 nm) pit membranes (Fig. S2), and there is no evidence for large (> 50 nm) pore size differences among species (Zhang *et al.*, 2020). So why then is xylem embolism resistance, which is frequently quantified as the xylem water potential corresponding to 50% loss of the maximum hydraulic conductivity (*P*_50_, MPa), so variable within angiosperms (Choat *et al.*, 2012)?

**Figure 1.**
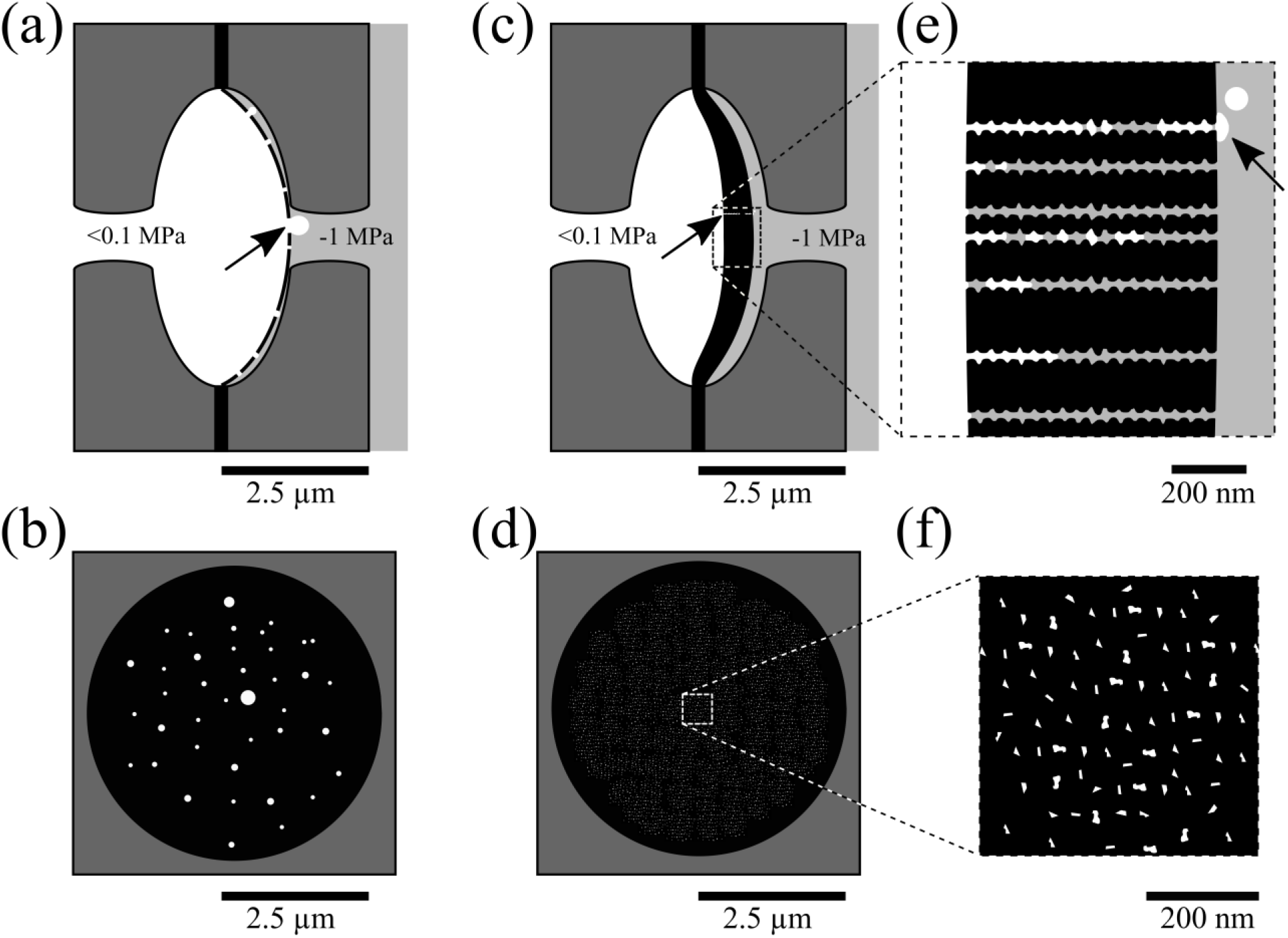
Drawings illustrating a mainly two-dimensional (a, b) and three-dimensional (c, d, e, f) concept of angiosperm pit membranes and air-seeding under aspiration. The upper images (a, c, e) show a longitudinal view, while the bottom ones (b, d, f) represent frontal views. Large, cylindrical pores with circular cross-sections occur in a pit membrane, with no defined thickness, and the largest pore triggers air-seeding (arrows in a, c, e). Pores in a 670 nm thick pit membrane that is composed of multiple layers of cellulose fibrillar aggregates show multiple pore constrictions, which greatly reduces the size of the narrowest constriction within a pore (c, f). A magnified view is shown in e and f, with seven pores in e), 18 pore constrictions per idealised pore cylinder. White colour = gas; bright grey = xylem sap; black = solid phase of the primary cell wall, middle lamella or pit membrane, dark grey = secondary cell wall.

Angiosperm species with thick pit membranes were found to be more resistant to drought-induced embolism than species with thin pit membranes (Jansen *et al.*, 2009; Li *et al.*, 2016). This functional link between *T*_PM_ and *P*_50_ is valid at the interspecific and intraspecific level (Lens *et al.*, 2011; Plavcová & Hacke, 2012; Scholz *et al.*, 2013; Schuldt *et al.*, 2016). Variation in *T*_PM_ is mainly determined by the number of microfibril layers (*N*_L_), with thin pit membranes consisting of fewer microfibril layers than thick pit membranes. Note that *N*_L_ can be estimated by assuming that cellulose fibres have a diameter of about 20 nm (Pesacreta *et al.*, 2005), and 20 nm pore spaces between each layer based on gold perfusion experiments (Table 1; Zhang *et al.*, 2020). As such, pit membranes with a thickness between 140 and 1,180 nm (Jansen *et al.*, 2009; Li *et al.*, 2016) include between 4 and 30 layers. In our models, bottlenecks in a given pore are formed by the pore constrictions between cellulose fibres within a single layer. Therefore, the number of constrictions within a pore (*N*_C_) equals *N*_L_ (Table 1). Since it is unknown why thin pit membranes are more vulnerable to embolism than thick pit membranes (Jansen *et al.*, 2018), we explore the hypothesis that the likelihood of leaky pores is affected by *N*_L_, which would explain why *T*_PM_ is related to *P*_50_.

**Table 1.** Overview of pit membrane thickness values (*T*_PM_, nm) and their corresponding numbers of microfibril layers (*N*_L_) according to the shrinkage model of Zhang et al. (2019). Assuming a homogeneous distribution of cellulose fibres, which have a diameter of 20 nm and a distance of 20 nm from each other, *N*_L_ = (*T*_PM_ + 20) / 40.

| $T_{PM}$ [nm] | 140 | 300 | 460 | 620 | 780 | 940 | 1100 | 1260 |
| --- | --- | --- | --- | --- | --- | --- | --- | --- |
| $N_L$ | 4 | 8 | 12 | 16 | 20 | 24 | 28 | 32 |

**Table 2:**
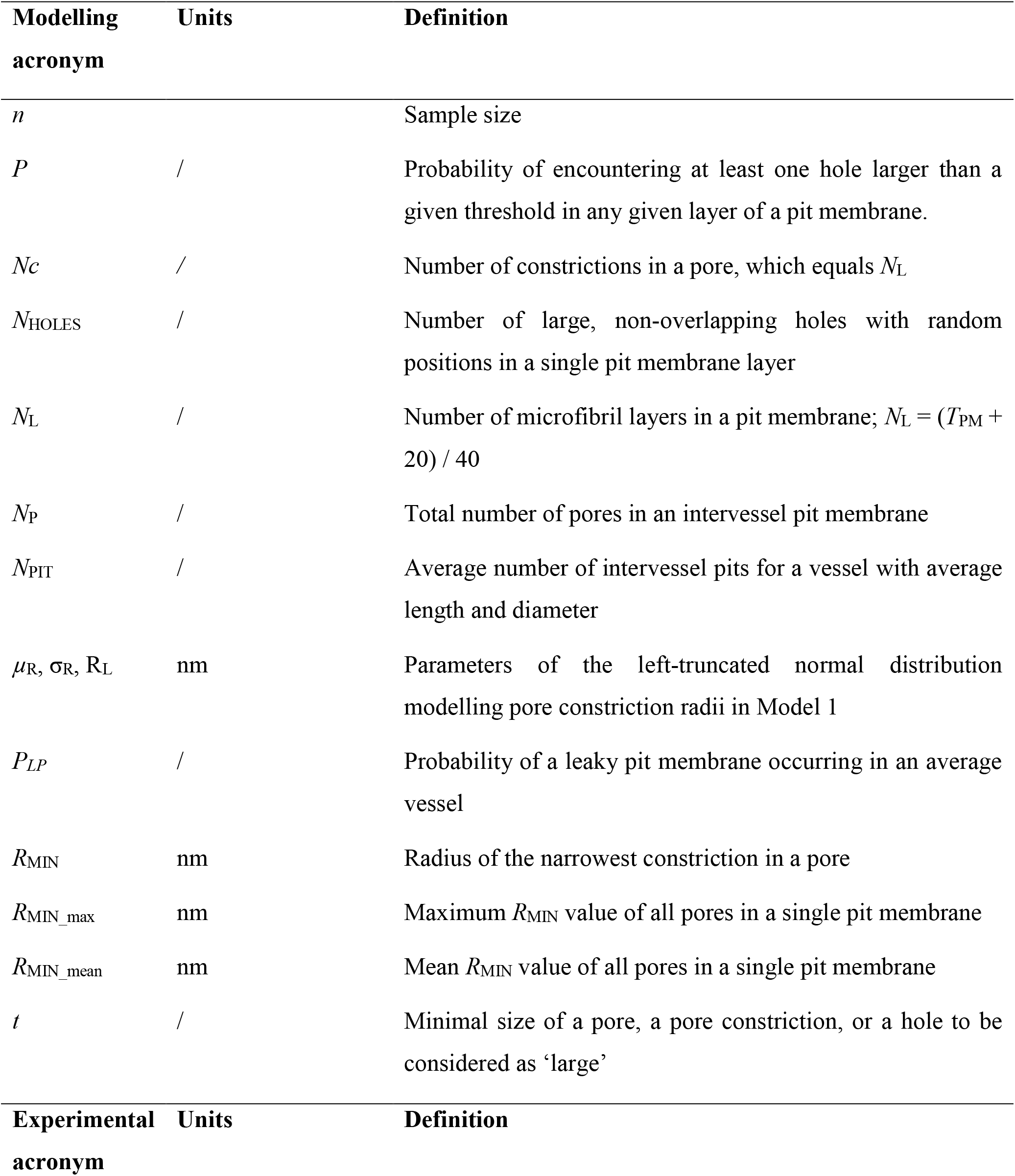
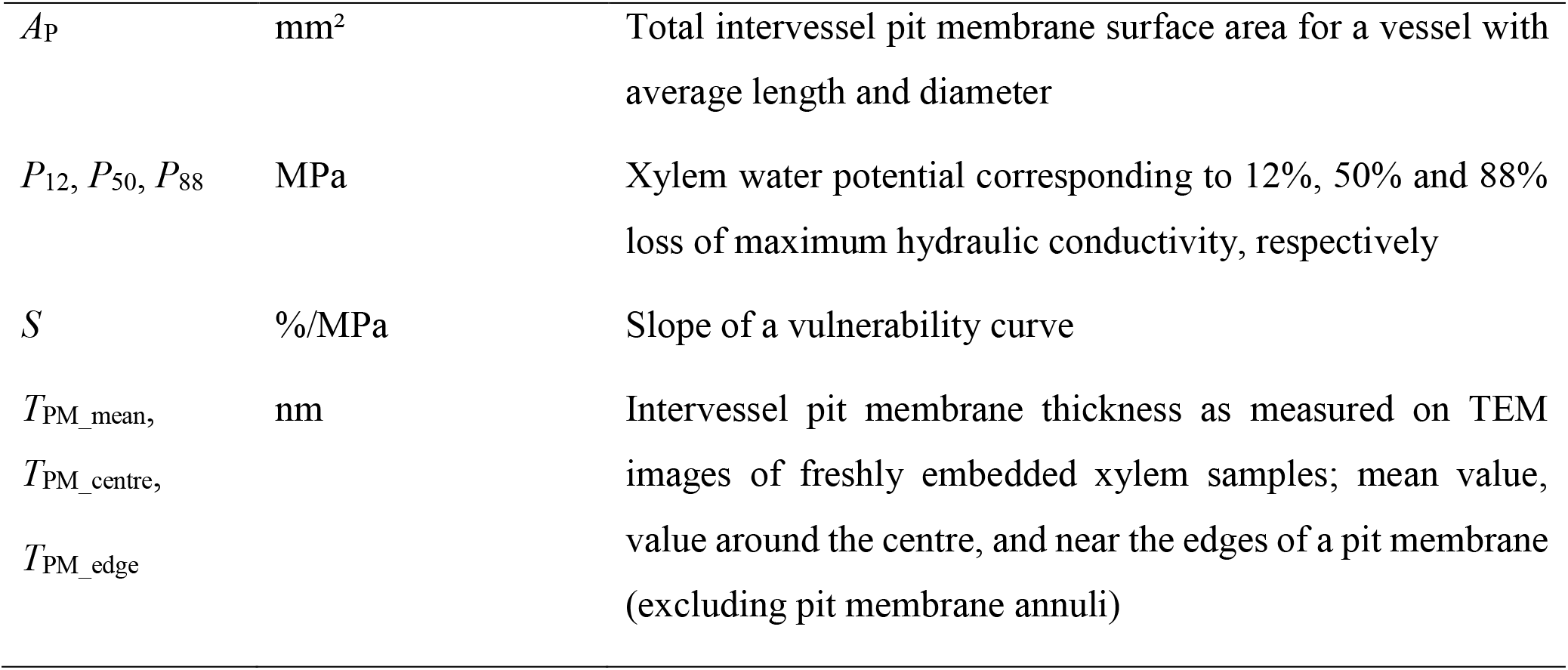
Overview of the abbreviations of modelling and experimental parameters used with reference to their units and definitions.

| Modelling<br>acronym | Units | Definition |
| --- | --- | --- |
| $n$ | | Sample size |
| $P$ | / | Probability of encountering at least one hole larger than a given threshold in any given layer of a pit membrane. |
| $N_C$ | / | Number of constrictions in a pore, which equals $N_L$ |
| $N_{\text{HOLES}}$ | / | Number of large, non-overlapping holes with random positions in a single pit membrane layer |
| $N_L$ | / | Number of microfibril layers in a pit membrane; $N_L = (T_{\text{PM}} + 20) / 40$ |
| $N_P$ | / | Total number of pores in an intervessel pit membrane |
| $N_{\text{PIT}}$ | / | Average number of intervessel pits for a vessel with average length and diameter |
| $\mu_R, \sigma_R, R_L$ | nm | Parameters of the left-truncated normal distribution modelling pore constriction radii in Model 1 |
| $P_{LP}$ | / | Probability of a leaky pit membrane occurring in an average vessel |
| $R_{\text{MIN}}$ | nm | Radius of the narrowest constriction in a pore |
| $R_{\text{MIN\_max}}$ | nm | Maximum $R_{\text{MIN}}$ value of all pores in a single pit membrane |
| $R_{\text{MIN\_mean}}$ | nm | Mean $R_{\text{MIN}}$ value of all pores in a single pit membrane |
| $t$ | / | Minimal size of a pore, a pore constriction, or a hole to be considered as ‘large’ |
| Experimental<br>acronym | Units | Definition |
| $A_P$ | mm<sup>2</sup> | Total intervessel pit membrane surface area for a vessel with average length and diameter |
| $P_{12}, P_{50}, P_{88}$ | MPa | Xylem water potential corresponding to 12%, 50% and 88% loss of maximum hydraulic conductivity, respectively |
| $S$ | %/MPa | Slope of a vulnerability curve |
| $T_{PM\_mean},$<br>$T_{PM\_centre},$<br>$T_{PM\_edge}$ | nm | Intervessel pit membrane thickness as measured on TEM images of freshly embedded xylem samples; mean value, value around the centre, and near the edges of a pit membrane (excluding pit membrane annuli) |

The mismatch between pore size estimations based on colloidal gold perfusion, and those calculated from vulnerability curves or air-seeding measurements, resulted in the hypothesis that a very small percentage of pit membranes might contain large, leaky pores (Choat *et al.*, 2003, 2004). These rare pit membrane pores are assumed to account for low air-seeding pressures (< 1 MPa). The idea of such leaky, rare pits was further enhanced when variation in *P*_50_ at an interspecific level was found to decrease with increasing pit membrane surface area in intervessel walls (Wheeler *et al.*, 2005). The “pit area hypothesis” (Sperry *et al.*, 2006), which was later termed “rare pit hypothesis”, provided a possible explanation for low air-seeding pressures, and relied on a largely two-dimensional interpretation of pit membranes (Hacke *et al.*, 2007; Christman *et al.*, 2009, 2012; Plavcová *et al.*, 2013). While the rare pit hypothesis follows a plausible concept that seems well supported by indirect evidence, it cannot be tested because the existence of a rare pit with a leaky pore cannot be observed directly, and is impossible to be verified from a statistical point of view. However, a three-dimensional modelling approach to estimate the likelihood of leaky pits is clearly lacking.

The number of layers in a pit membrane affects the size of the narrowest constriction within a pore that crosses the entire intervessel pit membrane (Fig. 1). Because the pressure difference required for air-seeding is determined by the radius of a meniscus in a pore, the most important dimension of a pore is its minimum diameter, i.e., the diameter of the narrowest bottleneck along the pore. We can think of this diameter as the “effective diameter” of the pore. The entry of an air-water meniscus or a bubble in a pit membrane is determined by the pore with the largest effective diameter within the pit membrane. Thus, air-seeding pressure and the minimum hydraulic resistance at the intervessel level is governed by the pore with the largest effective diameter in all pit membranes of a single vessel.

First, we hypothesise that the effective diameter of each pore becomes smaller with increasing *T*_PM_ and *N*_L_ (Hypothesis 1). This hypothesis is investigated at the individual pit membrane level based on a stochastic pit membrane model. Second, we hypothesise that model-based values of air-seeding pressure largely agree to embolism resistance measurements for a large number of species (Hypothesis 2). Third, we expect that the probability of having a leaky pit membrane is low at the whole vessel level, and affected by both *T*_PM_ (Li *et al.*, 2016), and the total number of intervessel pits per vessel (*N*_PIT_; Hypothesis 3) (Wheeler *et al.*, 2005). The second hypothesis is tested based on experimental data on embolism resistance, and anatomical measurements, while two further stochastic pit membrane models are developed to test the third hypothesis. Verifying these hypotheses should help us to better understand the functional link between embolism resistance and pit membrane ultrastructure.

## Materials and Methods

### Pit membrane modelling

To better understand the relationship between *T*_PM_ and *P*_50_, we developed three pit membrane models with different levels of complexity. For reasons of simplicity, we assumed the existence of more or less cylindrical pores, which govern transport phenomena. However, we modelled each pore as a three-dimensional object instead of a circular, flat opening (Sperry and Hacke, 2004; Mrad *et al.*, 2018). Following the multi-layered pit membrane model of Zhang *et al.* (2020), we assumed that each pore penetrates a fixed number of microfibril layers. Each of these layers induces a pore constriction of some random radius (Fig. 1e). An important property of each pore is its effective radius, i.e., the radius of the narrowest pore constriction within the entire pore (*R*_MIN_, nm). We were especially interested in how *R*_MIN_ was affected by *T*_PM_ (Hypothesis 1), how modelled air-seeding pressure based on pore constriction size related to measured embolism resistance (Hypothesis 2), and to what extent the likelihood of leaky pit membranes at the entire vessel level was affected by *T*_PM_ and/or *N*_PIT_ (Hypothesis 3).

Selected *T*_PM_ values spanned a range of 160 to 1,200 nm based on TEM (Jansen *et al.*, 2009; Li *et al.*, 2016). We assumed that the number of pore constrictions (*N*_C_) was equal to the number of microfibril layers *N*_L_, where *N*_L_ = [(*T*_PM_ +20) /40], and that the thickness of each layer corresponded to a single microfibril’s average diameter of 20 nm (Jansen *et al.*, 2009; Pesacreta *et al.*, 2005), with a distance of 20 nm between neighbouring layers. The latter seemed reasonable based on gold perfusion experiments (Choat *et al.*, 2003, 2004; Zhang *et al.*, 2017, 2020) and the fact that cellulose microfibrils are slightly negatively charged.

We developed three different models to investigate the relationship between *T*_PM_ and the probability of encountering at least one pore with *R*_MIN_ larger than a given threshold. While Model 1 was used to estimate pit membrane leakiness at the structural level of a single pit membrane, Models 2 and 3 considered leakiness at the vessel level.

### Model 1. Pore constrictions in single intervessel pit membranes

In this model (Fig. 2a), we assumed that a pit membrane comprised a fixed number of pores (*N*_P_), which were independent of each other, but we did not consider the location of pores within a pit membrane. Each pore was defined by a fixed number of pore constrictions (*N*_C_), given by the number of layers *N*_L_. The random radius of each pore constriction was modelled by a left-truncated normal distribution with parameters *R*_L_, *μ*_R_ and *σ*_R_, where *μ*_R_ and *σ*_R_ were mean and standard deviation of the untruncated normal distribution, and *R*_L_ was the lower bound for truncation. For a circular pit membrane with a diameter of 5 μm (estimated from *n* = 43 species, Jansen *et al.* 2009, 2011) and two different scenarios *R*_L_ = 2.5 nm, *μ*_R_ = 10 nm, *σ*_R_ = 7.5 nm (Scenario 1), and *R*_L_ = 2.5 nm, *μ*_R_ = 50 nm, *σ*_R_ = 40 nm (Scenario 2), we estimated an upper bound for the number of pores that might possibly fit into the membrane. The full specifications of the two scenarios were calculated as *N*_P_ = 12,000 (Scenario 1), and *N*_P_ = 1,100 (Scenario 2). While Scenario 1 was considered to be realistic based on gold perfusion experiments (Zhang *et al.*, 2020), Scenario 2 was taken as a conservative approach. For each pore, we considered *N*_C_-values between 4 and 32, and *T*_PM_-values between 140 and 1260 nm (Table 1). The random diameters of pore constrictions of a whole pit membrane was simulated ten times for the Scenarios 1 and 2, with *R*_MIN_ determined for each pore. Then, for each value of *N*_C_ we calculated the percentage of pores with *R*_MIN_ above a given threshold *t*, i.e., the percentage of leakiness for the modelled pit membrane. This threshold was chosen at *t* = *μ*_R_ + *σ*_R_, i.e., at 35 nm (Scenario 1) and 180 nm (Scenario 2). Furthermore, for each value of *N*_L_, the mean (*R*_MIN_mean_) and maximum values (*R*_MIN_max_) of the effective radii *R*_MIN_ obtained in the repeated simulation runs, were calculated.

**Figure 2.**
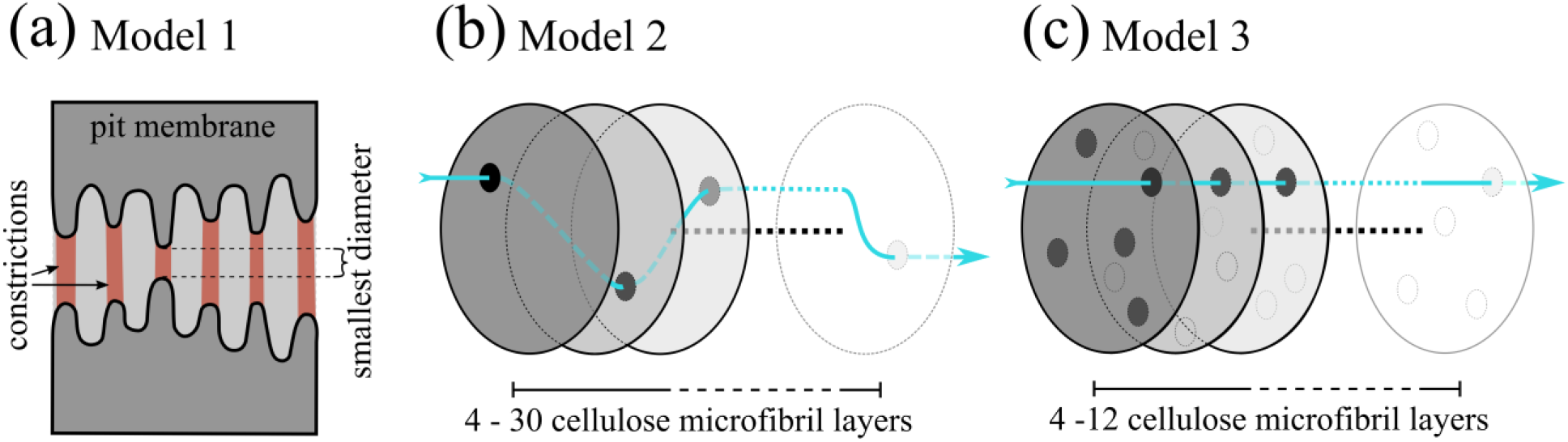
Three mathematical models to investigate the functional link between pit membrane thickness and effective diameters of pores. Model 1 (a) is based on a random number model to estimate the size of the narrowest constrictions of pores that traverse an entire pit membrane. This model was run ten times on a Scenario 1 with smaller and a Scenario 2 with larger pore constrictions for 12,000 or 1,100 pores per pit membrane, respectively, with 4 to 30 constrictions per pore in 140 to 1,200 nm thick pit membranes. Model 2 (b) examines the probability of large pores in 3,000 to 400,000 intervessel pit membranes within an entire vessel. Pit membranes included from four to 30 microfibril layers, assuming either a 25% or 50% chance of encountering a large hole in each layer. This model is independent of what we consider a large pore, and does not incorporate alignment of pore constrictions. Model 3 (c) evaluates the probability of encountering pores with a large effective radius at the vessel level (i.e. for 30,000 intervessel pits), with pit membranes consisting of 4 to 12 microfibril layers, assuming 5 or 10 pore constrictions of 200 nm per layer. Alignment of pore constrictions was included in Model 3 by simulating random locations of pore constrictions in each pit microfibril, and requiring minimal overlap between consecutive pore connections to create a pore. Different shades of grey represent various microfibril layers, and a hypothetical flow path is indicated by the blue lines in (b) and (c).

To compare the results obtained from Model 1 with experimental data on embolism resistance, the theoretical air-seeding pressure was calculated based on *R*_MIN_max_ and *R*_MIN_mean_. For this, a modified Young-Laplace equation was applied:

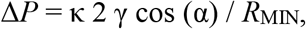

where Δ*P* was the pressure required to induce air-seeding, κ was a dimensionless pore shape correction factor which was assumed to be equal to 0.5 (Schenk *et al.* 2015), γ was the surface tension of xylem sap, α represented the contact angle of the gas-xylem sap interface with the solid cellulose microfibril and was assumed to be equal to zero, and *R*_MIN_ was the narrowest pore constriction radius. We assumed that γ was either 72 or 25 mN/m, which corresponded to the surface tension of pure water and was close to the bulk surface tension of xylem sap (Christensen-Dalsgaard and Tyree, 2014), and the equilibrium surface tension of xylem sap lipids based on dynamic surface tension (Schenk *et al.*, 2017; Yang *et al.*, 2020).

### Models 2 and 3. Leaky pit membranes at the vessel level

Model 2 investigated the occurrence of leaky pit membranes at the vessel level. We first calculated an upper bound for the probability that a leaky pore would run through an entire intervessel pit membrane (Fig. 2b). The minimum requirement needed for a leaky pore was that there existed at least one hole with a radius larger than a given threshold *t* in each layer. The term hole was used as a substitute for constriction as its diameter might even exceed the length of the pore. We did not account for proper alignment of the holes in Model 2. The probability that at least one large hole existed in each layer, was an upper bound for the probability of encountering at least one pore through the whole membrane with an effective radius larger than *t*. We assumed that the probability *P* of encountering a large hole in any given layer is independent of the occurrence of this event in other layers, and constant across all layers. In particular, we assumed that a probability of *P* = 0.25 represents a safe scenario, whereas *P* = 0.50 represents a risky scenario. The probability of encountering at least one leaky pore through the whole membrane was given by 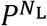. At the entire vessel level, an upper bound for the probability of having a leaky pit membrane (*P*_LP_) was given by

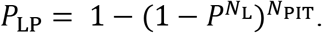

We estimated *N*_PIT_ based on the total pit membrane surface area per vessel (*A*_P_) of 72 species using original data and literature data (Fig. S1; Wheeler *et al.*, 2005; Jansen *et al.*, 2011; Lens *et al.*, 2011; Nardini *et al.*, 2012; Scholz *et al.*, 2013; Klepsch *et al.*, 2016). Values of *N*_PIT_ were not normally distributed (Shapiro-Wilk test, W = 0.84, *p* < 0.0001, *n* = 65, excluding outliers), and were varying asymmetrically from 510 to 370,755, with a median of 14,188 (IQR: 31,618, Q_1_: 4,651 *N*_PIT_, Q_3_: 36,269, *n* = 72 species). Since only six species showed *N*_PIT_-values above 83,068 (Q_3_ + 1.5 * IQR), we took *N*_PIT_ = 80,000 as a representative maximum value in our graphs, although we investigated the occurrence of leaky pits up to *N*_PIT_ of 400,000.

Since we did not consider in Model 2 the alignment of holes within successive layers, we effectively assumed that gas could spread between layers of the membrane and then entered through the next large holes of the following layer. Although it is currently unknown whether such an alignment of holes would be required for air-seeding, we incorporated the location of holes in pit membrane layers in Model 3. More precisely, we modelled pit membranes as a stack of *N*_L_ circular layers of cellulose material, with diameter *D*_P_ (= 5 μm), and no gap between two adjacent layers. Each layer was perforated by randomly located holes. Since we were interested in pores with a minimum radius larger than a given threshold *t*, holes smaller than this threshold were ignored. Thus, in each layer, we randomly placed a fixed number of large, non-overlapping holes (*N*_HOLES_). For simplicity, we chose the threshold *t* as radius for all these holes. Then, we determined the pores that crossed the whole pit membrane. A pore did only traverse all layers if there existed a sequence of holes such that for each pair of adjacent layers, the holes were properly aligned and overlapping (Fig. 2c). Holes without minimal overlapping with holes in adjacent layers were assumed to form dead-ends and ignored. The locations of holes within and across layers were simulated stepwise and repeated 10^6^ times for pit membranes with *N*_L_-values between 4 and 12 (Table 1). Details of the implementation are given in the Supporting Information (Method S1). For simplicity, we ignored incomplete overlap of adjacent holes, which could lead to a reduced minimum radius of the resulting pore. Using this model, we modelled a large number of membranes for *t* (hole threshold radius) = 200 nm, *D*_P_ = 5 μm, *N*_L_ = 4 to 12, and *N*_HOLES_ = 5 or 10. These values of *R*_MIN_ and *N*_HOLES_ were selected to make pit membranes leakier than available evidence suggests, to increase the likelihood of overlapping holes, and to avoid underestimating leakiness. For each scenario, we estimated the probability that at least one leaky pore with *R*_MIN_ ≥ 200 nm crossed an entire pit membrane. Finally, we estimated the probability of leaky pit membranes with at least one large pore for a vessel with 30,000 intervessel pits (*N*_PIT_), which was well above the median *N*_PIT_ of 14,188 intervessel pits (Fig. S1).

### Experimental work

The three models were complemented by experimental data on embolism resistance (*n* = 31 species), *T*_PM_ measurements at the centre (*T*_PM_centre_) and near the edges (*T*_PM_edge_) (*n* = 31 species), and the total intervessel pit membrane area per average vessel (*A*_P_, *n* = 20 species). The methods applied to obtain these data include well-established, previously published protocols (Wheeler *et al.*, 2005; Sperry *et al.*, 2006; Schuldt et al. 2016; Zhang *et al.* 2020; Kotowska *et al.*, 2020), and are described in detail in the Supporting Information (Method S2). All data include original measurements, except for data retrieved from literature for embolism resistance of five species, and for *A*_P_ values of four species.

### Statistics and data processing

Data processing, simulations and statistical analyses were performed using Excel, R, and Matlab. Shapiro-Wilk Tests were applied to test for normal distribution. Pearson’s Correlation Coefficient were used to test for linear correlation. Basic linear and non-linear regressions were fitted to test whether *P*_12_, *P*_50_, *P*_88_, and the slope of vulnerability curves (*S*) are related to *T*_PM_ or *A*_P_, and could be estimated. Only significant regressions with the highest R^2^ were considered. For each of the 31 species studied, we estimated air-seeding pressures by integrating their modelled *R*_MIN_mean_ and *R*_MIN_max_, based on *T*_PM_, into the equations of the relation between *T*_PM_ and air-seeding pressure of Model 1. This approach allowed us to compare estimated air-seeding with experimental values of *P*_12_ and *P*_50_. Model 3 was simulated using R (Method S1).

## Results

### How likely are large pores in a pit membrane for a wide range of pit membrane thicknesses?

Average values of *R*_MIN_ (*R*_MIN_mean_) are very low in Scenario 1 of Model 1, with values below 9 nm for pit membranes with 150 to 1,150 nm in thicknesses (Fig. 3a). The size of *R*_MIN_ declines considerably with increasing *T*_PM_, and the largest ones (*R*_MIN_max_) decrease from ca. 40.7 ± 2.7 to 12 ± 1.1nm (Fig. 3a). *R*_MIN_max_-values are at least 2.4 times and up to 4.9 times larger than the *R*_MIN_mean_-values, decreasing with *T*_PM_ (Fig. 3a). The likelihood of having an effective diameter ≥ 35 nm approaches zero (0.00005 ± 0.00009, Fig. 3b) when *T*_PM_ is > 220 nm, or *N*_L_ ≥ 6, thus only occurs in 0.2 out of 12,000 pores.

**Figure 3.**
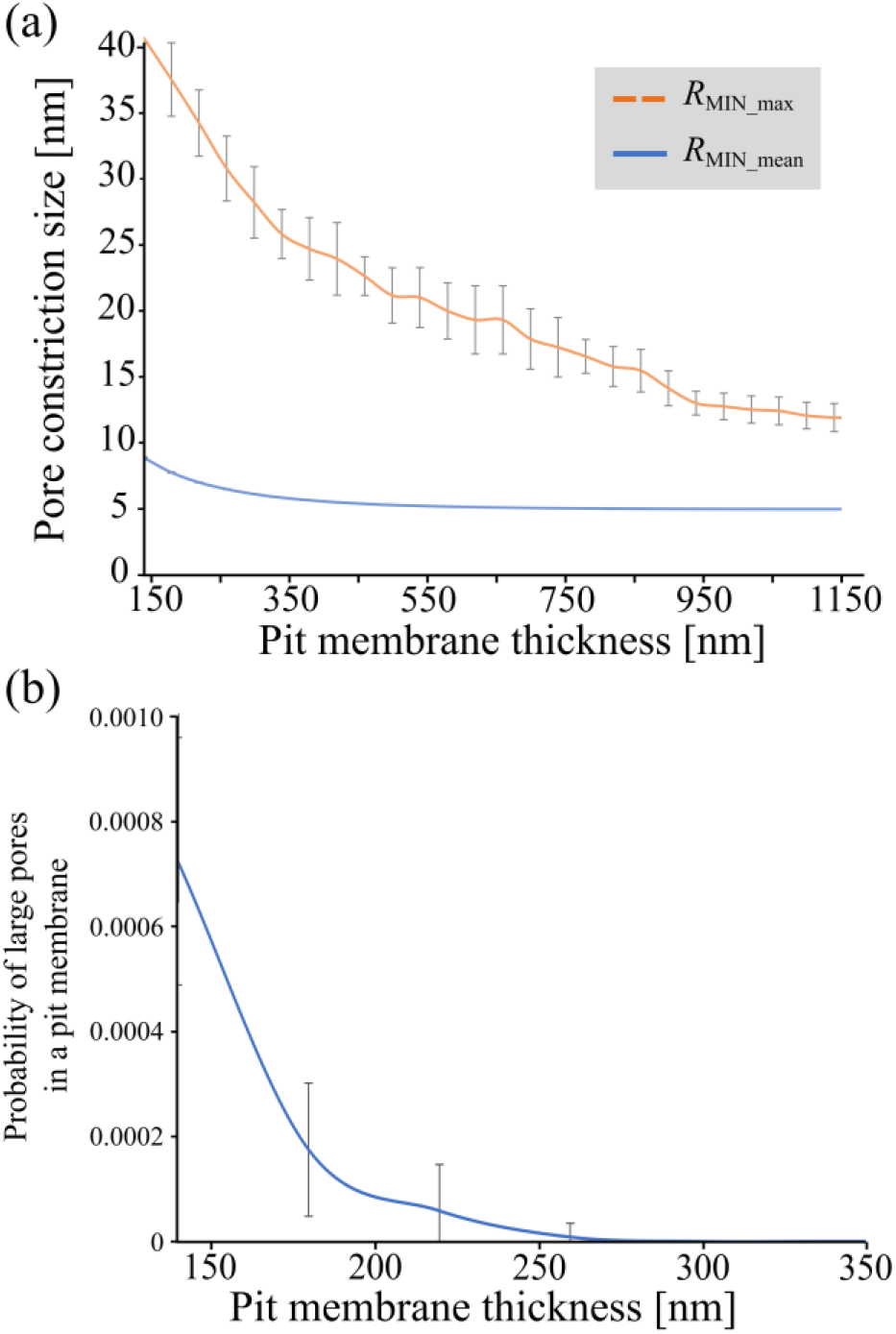
Results of Scenario 1 of Model 1, showing the pit membrane thickness plotted versus the pore constriction diameter based on Model 1 (a), and the likelihood of a relatively large *R*_MIN_max_ (≥ 35 nm) within a pit membrane (b), which decreased exponentially from 0.0008 ± 0.0002 to values approaching zero with increasing pit membrane thickness. A random number model was used, with the mean pore constriction size set to 20 ± 15 nm, and a minimum size of 5 nm. Pore constriction sizes were determined ten times for 12,000 simulated pores, corresponding to an average sized pit membrane.

For Scenario 2 of Model 1, a similar decline of *R*_MIN_ with increasing *T*_PM_ is found (Fig. S3), but with steeper declining likelihood values for large pores with *T*_PM_. For a *T*_PM_ of 220 nm the likelihood of containing a large pore (defined in Scenario 2 of Model 1 as ≥ 180 nm in diameter) is nearly zero.

### How does pit membrane thickness relate to measured embolism resistance?

The values of *T*_PM_mean_ vary from 165 nm (± 18 SD) for *Tilia platyphyllos* to 610 nm (± 79 SD) for *Olea europaea*, and the median of *T*_PM_ is equal to 270 nm (*n* = 31 species studied; Table S1). The value of *T*_PM_centre_ is always larger than the value of *T*_PM_edge_, with an average difference of 105 nm, varying from 2.1 nm (*Tilia platyphyllos*) to 297 nm (*Olea europaea*), and increasing with *T*_PM_.

*P*_50_-values are strongly related to the values of *T*_PM_centre_ (Table 3; Fig. 4c), with a logarithmic regression showing an R^2^-value of 0.57 (F(2, 29) = 32.0, *p* < 0.001). An outlier in the *T*_PM_ vs. *P*_50_ relationship includes *Corylus avellana*, which shows considerably high *T*_PM_-values of ca. 400 nm for a *P*_50_–value of −2.02 MPa. Slightly lower correlations are found between the *T*_PM_centre_ and *P*_12_ (F(2, 29) = 24.4, *p* < 0.001, R^2^ = 0.457), and between *T*_PM_centre_ and *P*_88_ (F(2, 29) = 34.2, *p* < 0.001, R^2^ = 0.541; Table 3). Thus, the *T*_PM_centre_-values show a stronger relationship to embolism resistance than *T*_PM_mean_ and *T*_PM_edge_. The average intervessel pit membrane surface area per vessel (*A*_P_, Table S1) shows much lower correlations to *P*_50_, *P*_12_ and *P*_88_ than all *T*_PM_ traits, with the strongest correlation between *A*_P_ and *P*_12_ (F(2, 18) = 7.75, *p* < 0.05, R^2^ = 0.301; Table 3).

**Table 3.** Overview of the r- and R²-values between pit anatomical characteristics and embolism resistance. Anatomical measurements include mean values of the intervessel pit membrane thickness (*T*_PM_mean_), central measurements (*T*_PM_centre_), those near the pit membrane annulus (*T*_PM_edge_), and the total intervessel pit membrane area per vessel (*A*_P_). Embolism resistance has been quantified as xylem water potential values corresponding to 12% (*P*_12_), 50% (*P*_50_), and 88% (*P*_88_) loss of the maximum hydraulic conductivity based on vulnerability curves. The estimation of air seeding pressure is either based on the largest value of *R*_MIN_ across all pores of a membrane (Air seeding *RMIN_max*) or the mean value of *R*_MIN_ across all pores of a membrane (Air seeding *R*_MIN_mean_), using a modified Young-Laplace equation. The regression models that show the strongest relation are given here. Logarithmic regression^1^; power regression^2^; Pearson Coefficient Correlation^3^; p-values: < 0.05 = *, < 0.01 = **, < 0.001***.

| | $P_{12}$ | $P_{50}$ | $P_{88}$ | $S$ | $T_{PM\_mean}$<br>SD |
| --- | --- | --- | --- | --- | --- |
| $T_{PM\_centre}$ | 0.457 ***<sup>1</sup> | 0.571 ***<sup>1</sup> | 0.541 ***<sup>1</sup> | 0.485 ***<sup>2</sup> | NA |
| $T_{PM\_mean}$ | 0.439 ***<sup>1</sup> | 0.556 ***<sup>1</sup> | 0.525 ***<sup>1</sup> | 0.477 ***<sup>2</sup> | 0.759 ***<sup>3</sup> |
| $T_{PM\_edge}$ | 0.306 **<sup>1</sup> | 0.410 ***<sup>1</sup> | 0.392 ***<sup>1</sup> | 0.340 ***<sup>2</sup> | NA |
| $A_P$ | 0.301 *<sup>1</sup> | 0.252 *<sup>1</sup> | 0.221 *<sup>1</sup> | 0.103 ***<sup>2</sup> | NA |
| **Air seeding**<br>$R_{MIN\_max}$ | 0.636 ***<sup>3</sup> | 0.732 ***<sup>3</sup> | NA | NA | NA |
| **Air seeding**<br>$R_{MIN\_mean}$ | 0.670 ***<sup>3</sup> | 0.739 ***<sup>3</sup> | NA | NA | NA |

**Figure 4.**
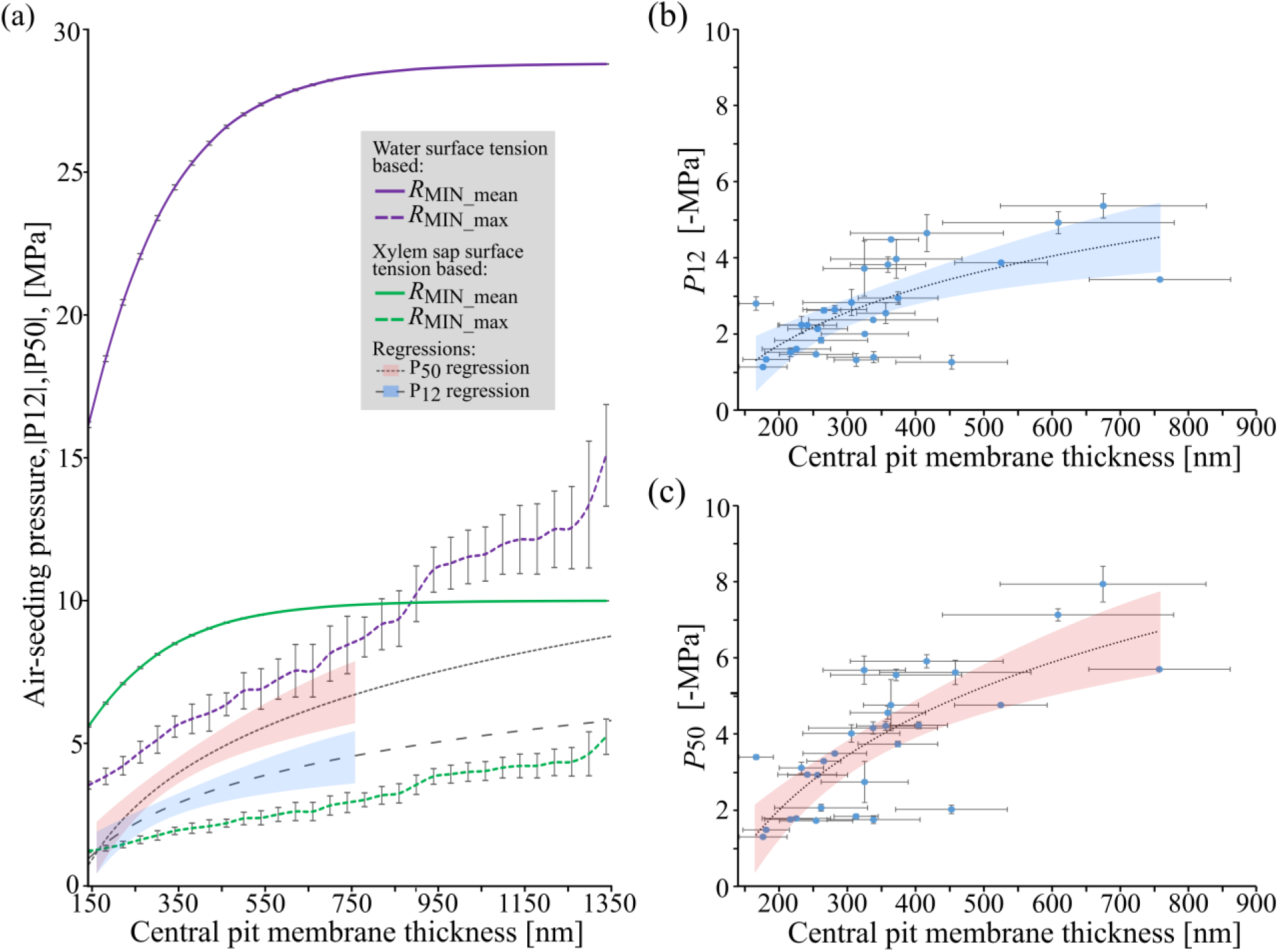
The relationship between pit membrane thickness and the modelled air-seeding pressure (a), measured *P*_12_ values (b), and measured *P*_50_ values of 31 angiosperm angiosperms (c). Modelled air-seeding pressures were based on the largest value of *R*_MIN_max_ (a; dotted green and purple lines), and *R*_MIN__min across all pores of a membrane (a; solid green and purple lines) for pit membranes with a wide range of thicknesses according to Model 1. We applied a modified Young-Laplace equation to obtain air-seeding pressure values for both 72 mN/m (purple lines) and 25 mN/m (green lines) surface tension (a). *P*_12_ and *P*_50_ values based on a flow-centrifuge method and microCT measurements plotted against central pit membrane thickness (*T*_PM_) measurements. *T*_PM_ was based on TEM of 31 plant species. Logarithmic regression line in dark grey (dashed) and corresponding confidence intervals (*P*_12_ = blue, *P*_50_ = red). In a) theses regression lines and confidence intervals are incorporated (*P*_12_ regression = wider dotted dark grey line, blue; *P*_50_ regression = denser dotted dark grey line, red).

*T*_PM_mean_ shows a linear relationship with *T*_PM_mean_ SD, with larger variation in thick than in thin pit membranes (Pearson’s Correlation Coefficient, r (29) = 0.759, *p* < 0.00001). Furthermore, we find a power regression with an R^2^–value of 0.477 between the slope of vulnerability curves (*S*) and *T*_PM_mean_ (F(2, 29) = 88.4, p < .001, R^2^ = 0.477; Table 3), with decreasing *S* being associated with increasing *T*_PM_mean_. There is a weaker relation between *S* and *T*_PM_edge_, and a slightly stronger relation to *T*_PM_centre_ than *T*_PM_mean_ (Table 3).

### Does modelled air-seeding correspond to measured embolism resistance for a wide range of pit membrane thicknesses?

There are clear differences in the estimated air-seeding pressures, depending on the surface tension, and whether the maximum or mean *R*_MIN_-values are considered (Fig. 4). For a surface tension of 72 mN/m, estimated air-seeding pressures, which theoretically correspond to *P*_12_, are much higher than the *P*_12_ values measured, and even higher than *P*_50_ measurements. Regression lines of the *T*_PM_-*P*_50_ and *T*_PM_-*P*_12_ relationship, however, fall well within the estimated air-seeding pressures when a surface tension of 25 mN/m (green lines in Fig. 4a) is considered. Although absolute values of modelled and measured air-seeding (*P*_12_) and embolism resistance pressures (*P*_50_) do not match (Fig. 4a, 5), they are significantly related to each other (Pearson’s Correlation Coefficient, *P*_12_ to *R*_MIN_mean_ and *R*_MIN_max_: r (29) = 0.67 and r (29) = 0.636, *p* ≪ 0.01; *P*_50_ to *R*_MIN_mean_ and *R*_MIN_max_: r (29) = 0.739 and r (29) = 0.732, *p* < 0.00001; Table 3, Fig. 5). When *R*_MIN_max_ is considered, estimated air-seeding pressures show a small range, with about 1.2 MPa for a *T*_PM_ of 140 nm and up to 2.7 MPa for a *T*_PM_ of 758 nm (Fig. 5b), which underestimates embolism resistance (Fig. 4a, 5a, 5b). Much higher air-seeding pressures between 5.6 and 10 MPa are obtained for estimations based on *R*_MIN_mean_, overestimating embolism resistance (Fig. 4a, 5c, d). There is a clear upper limit of air-seeding pressure for *R*_MIN_mean_ around ca. 10 MPa, which is achieved for pit membranes with thicknesses ≥ 600 nm.

**Figure 5.**
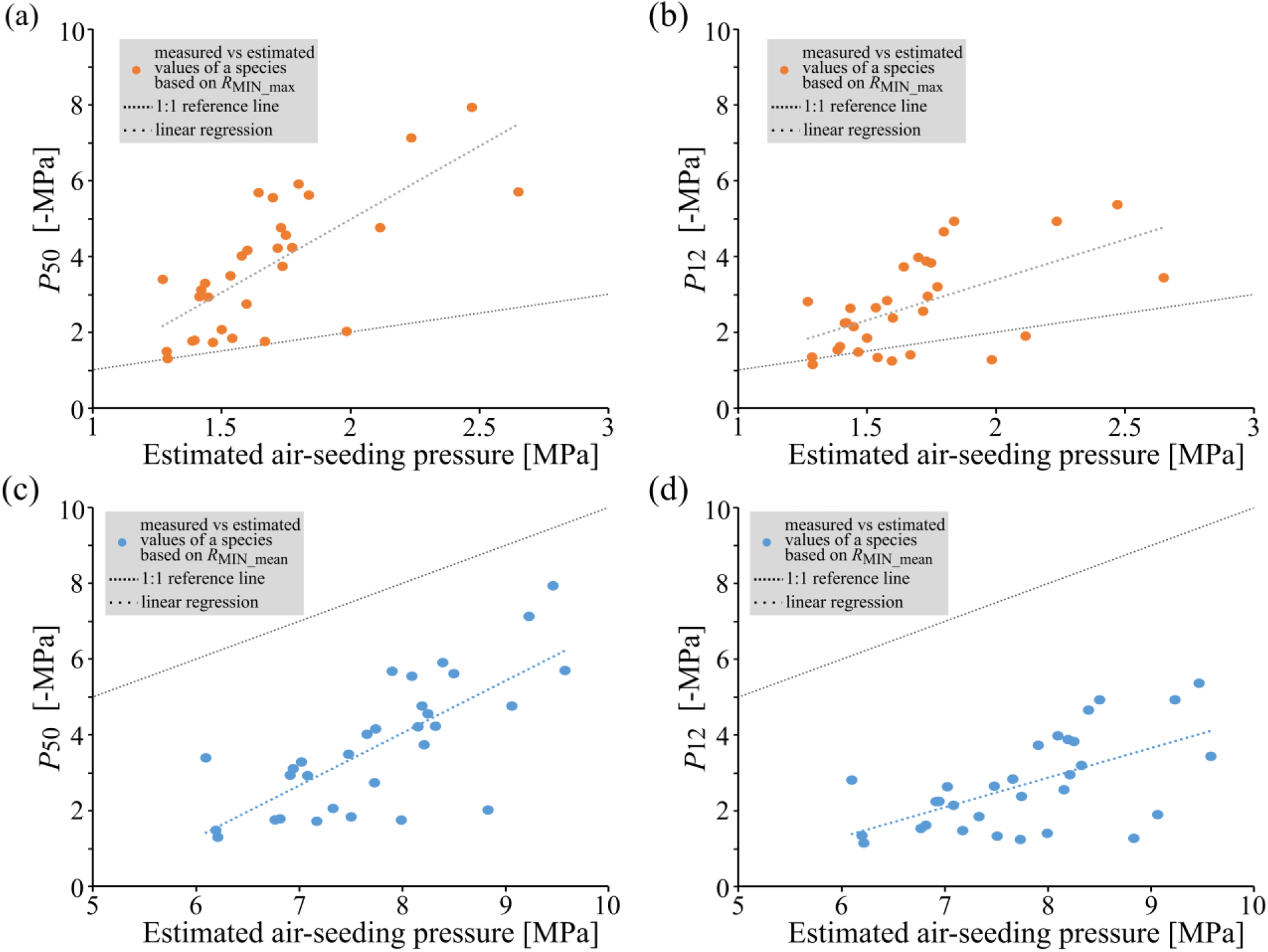
Modelled air-seeding pressure based on R_MIN_max_ (a, b) and R_MIN_mean_ (c, d) following Scenario 1 of Model 1 versus measured *P*_50_ (a, c) and *P*_12_ (b, d) for 31 angiosperm species.

Modelled air-seeding pressures based on *R*_MIN_max_ are similar but typically lower than the experimental values (Fig. 5a, b). Estimated air-seeding pressures based on *R*_MIN_max_ are especially close to measured embolism resistance for various species with not very negative *P*_12_- and *P*_50_-values (Fig. 5a, b), while estimated air-seeding pressures based on *R*_MIN_mean_ are much higher than *P*_12_ and *P*_50_ measurements (Fig. 5c, d).

### How likely are leaky intervessel pit membranes at the vessel level?

Based on Model 2, the probability of having a leaky pit membrane in a vessel decreases exponentially with increasing *T*_PM_ (Fig. 6, Fig. S4). For a fixed *T*_PM_, the slope of the relationship between *N*_PIT_ and the probability of a leaky pore strongly depends on *T*_PM_ (Fig. S5): steep, exponential slopes are found for thin pit membranes, while low, more linear slopes are found for thick pit membranes. Therefore, *T*_PM_mean_ and *N*_PIT_ affect the likelihood of large effective pore radii differently, with *N*_PIT_ having an unequal effect on the likelihood of having leaky pit membranes.

**Figure 6.**
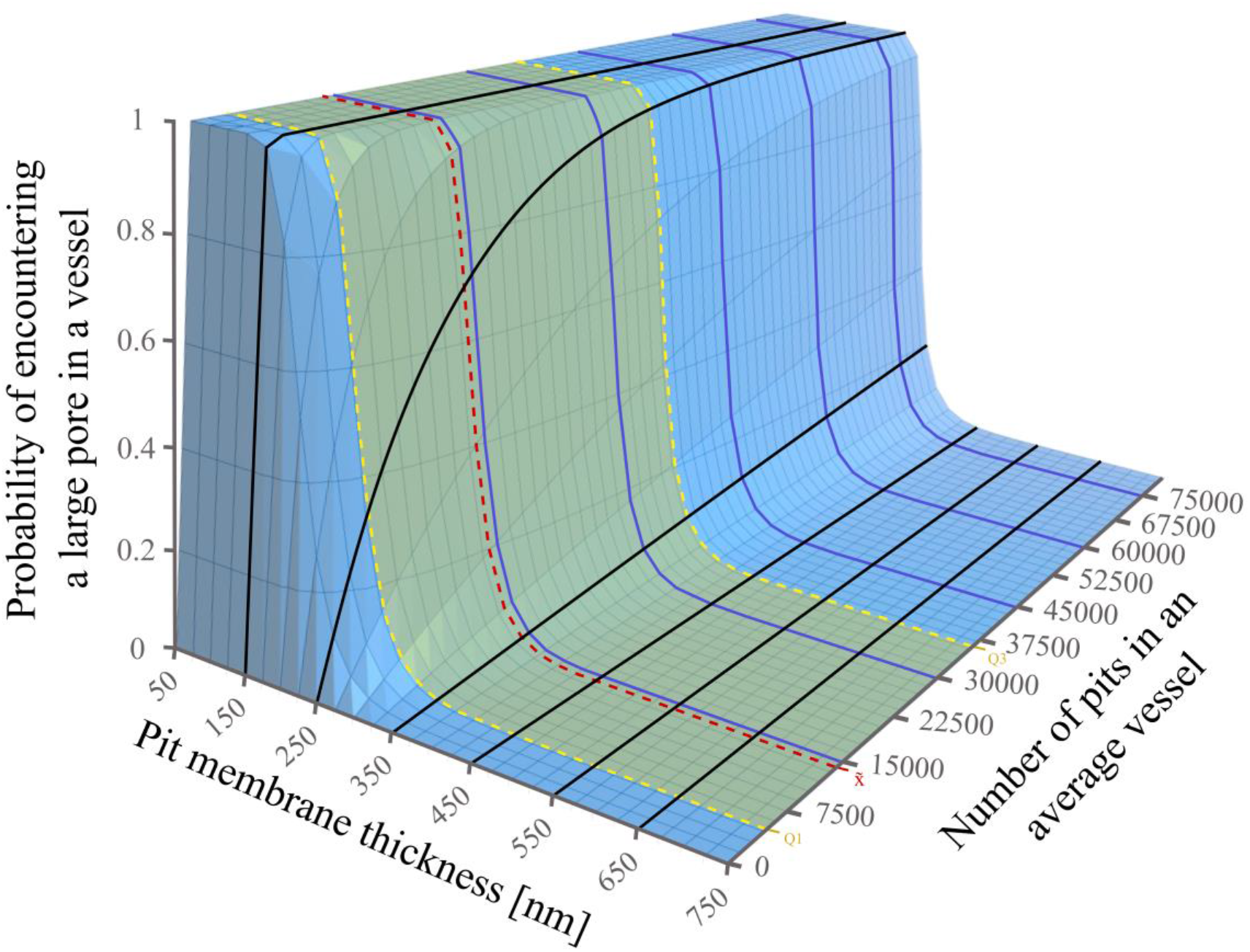
The probability of encountering at least one pore with large effective diameters in intervessel pit membranes for an entire vessel decreases with increasing pit membrane thickness (blue lines), but increases with increasing number of pits (black lines) according to Model 2. The likelihood of having a large hole within a single microfibril layer was assumed to be ≤ 0.25. This model did not consider the actual size of the pore constriction and ignored whether or not a hole was aligned with other holes in adjacent membrane layers. The green area indicates where most angiosperm species occur based on the number of intervessel pits per vessel, with the median (red dotted line), and the first and third quartile (yellow dotted line).

For the 0.5 likelihood assumption (Fig. S4, S5b), vessels with 820 nm thick pit membranes reach a likelihood of having a leaky pit membrane below 0.20, even in vessels with 400,000 intervessel pits, which means that not even every fifth vessel would have a leaky pit.

For the 0.25 likelihood of Model 2 (Fig. 6, S5a), an exponential change is found for *T*_PM_mean_-values between 200 and 300 nm, while little or no effect is seen for *T*_PM_mean_-values below 200 nm and above 350 nm. The high and low probability plateaus in the three-dimensional graphs of Model 2 (Fig. 6, S4) suggest the existence of a thin and a thick *T*_PM_-range that typically results in leaky or very safe, non-leaky vessels, respectively, independent of *N*_PIT_. At the exponential phase of the three-dimensional graph in Fig. 6, an increase in *N*_PIT_ from 3,000 to 70,000 (i.e. a 23-fold increase) is equivalent to adding about five additional microfibril layers to a pit membrane (i.e. an increase in *T*_PM_ of 180 nm). Critical *T*_PM_-values are higher for the 0.5 likelihood of Model 2 (Fig. S5b, S4), with the largest effect of *N*_PIT_ for pit membranes between 500 and 700 nm.

The results obtained from Model 3 show that the modelled probability of leaky pit membranes in a vessel with 30,000 intervessel pits (*N*_PIT_) decreases exponentially from 0.045 for 140 nm thick pit membranes to < 0.01 for *T*_PM_-values above 180 nm (Fig. 7). Assuming 5 or 10 holes per microfibril layer (*N*_HOLES_), less than one out of 30,000 pits has a large pore for *T*_PM_-values above 220 nm and 340 nm, respectively. Therefore, 220 nm thick pit membranes with a *N*_HOLES_-value of 5 have a similar safety as 340 nm thick pit membranes with an *N*_HOLES_-value of 10.

**Figure 7.**
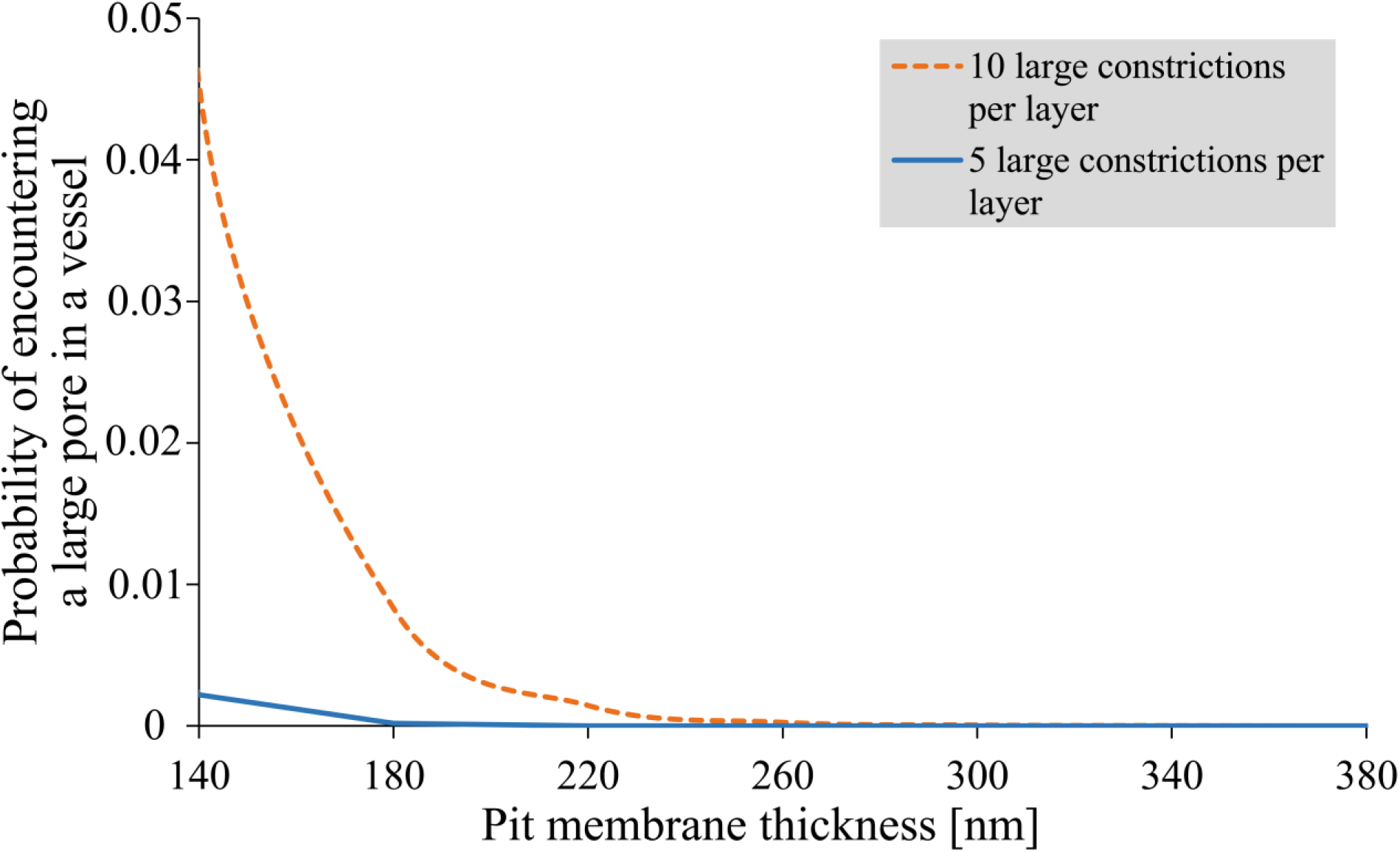
Output of Model 3, which assumed that intervessel pit membranes have a diameter of 5 μm, thicknesses between 140 nm and 460 nm, and an estimated number of 30,000 pits per vessel. Minimal overlapping of holes was required to obtain a pore through the whole membrane. The probability of pores with effective radius larger than a given threshold decreased exponentially in pit membranes of 140 nm to 260 nm in thickness when 10 holes of 200 nm in diameter occurred per pit membrane layer (orange line). Assuming 5 pore constrictions per layer (blue line) showed a very low likelihood of large pores, even for 140 nm thick pit membranes.

## Discussion

The results described above indicate that the chance of having large pores in pit membranes decreases strongly with the number of constrictions, and therefore *T*_PM_ (Hypothesis 1). This finding is independent of the actual size of pore constrictions, and supported by a strong relation between embolism resistance and *T*_PM_ (Jansen *et al.*, 2009, 2018; Lens *et al.*, 2011; Scholz *et al.*, 2013; Schuldt *et al.*, 2016; Li *et al.*, 2016). Modelled air-seeding values are significantly related to measured embolism resistance (Hypothesis 2), although they differ in absolute values. There is a good agreement when the dynamic surface tension of xylem sap is taken into account (Yang *et al.*, 2020), but embolism spreading does not seem to represent a function of pore constriction size (*R*_MIN_max_ and *R*_MIN_mean_) only. Our results also suggest that the likelihood of having a leaky pit membrane within a vessel is extremely low (Hypothesis 3), and mainly determined by *T*_PM_. Overall, pore constrictions provide a mechanistic explanation why embolism resistance is correlated with *T*_PM_, and why pit membranes provide hydraulic safety to angiosperm xylem.

### The most narrow pore constriction becomes strongly reduced in size with increasing pit membrane thickness

The three models developed show a negative correlation between the simulated pore sizes and *T*_PM_, which is reflected in a low probability of large pores, both at the level of an individual pit membrane and an entire vessel. Based on Model 1, the chance of having a large pore in a pit membrane thicker than 180 nm is close to zero. Interestingly, the thinnest pit membranes measured in this study (ca. 165 to 180 nm) are likely to represent a lower limit for *T*_PM_, since earlier records of *T*_PM_ below 150 nm (Jansen *et al.*, 2009; Li *et al.*, 2016) are likely artefacts due to shrinkage (Zhang *et al.*, 2017, 2018, 2020; Kotowska *et al.*, 2020). Thus, angiosperm pit membranes seem to have at least four or five layers of cellulose microfibrils and pore constrictions, which keeps the number of large pores very low for most species. There is a clear conceptual relationship between the thickness of a fibrous porous medium, and the size of the narrowest pore constriction as also seen for non-woven, fibrous geotextiles that differ in thickness (Aydilek *et al.*, 2007).

Model 2 suggests that the probability of encountering large pores in intervessel walls follows an exponential pattern over a fairly narrow range of *T*_PM_, with critical *T*_PM_-values between 200 to 300 nm and 500 to 700 nm for a 0.25- and 0.50-likelihood, respectively, of having at least one hole larger than *t* within a single microfibril layer. Although this likelihood cannot be accurately determined due to our limited understanding of air-seeding, we believe that a realistic likelihood would probably lay around 0.25, with 0.50 being too conservative. This assumption is supported by the steeper increase in embolism resistance within the lower *T*_PM_-range between 140 to 340 nm than in the higher *T*_PM_-range, and by the probabilities of large pores in pit membranes approaching zero for *T*_PM_ > 250 nm in Model 1 and 3. We applied a logarithmic regression between *P*_50_, *P*_12_ and *T*_PM_ (Fig. 4b, 4c), unlike a linear scaling that was previously suggested (Lens *et al.*, 2011; Li *et al.*, 2016). Interestingly, this logarithmic regression has *P*_50_-values approaching 10 MPa for a *T*_PM_ of > 1,000 nm (Fig. 4a), which corresponds to the upper physical limit of both xylem water potential and the maximum *T*_PM_–value measured (Vilagrosa *et al.*, 2003; Jansen *et al.*, 2009; Kanduč *et al.*, 2020).

Although we do not know whether alignment of holes across different layers is required for mass flow of air across a pit membrane, misalignment of holes could reduce the likelihood of having a leaky pit membrane. There is a very low chance of having a single, large pore in a vessel with 30,000 intervessel pit membranes having a *T*_PM_–value of 200 nm or more, even if extremely large holes with a diameter of 200 nm occur in a single microfibril layer (Model 2 and 3). It is possible that variation in *T*_PM_ within a vessel or within the vessel network provides additional chances of leakiness. Capturing this variation, however, is difficult because measuring pit membrane thickness may not be straightforward, for instance due to TEM preparation artefacts, aggregation of cellulose fibrils into larger aggregates, and seasonal shrinkage of pit membranes (Schmid & Machado, 1968; Sorek *et al.*, In press).

Aspiration or a mechanical pressure on intervessel pit membranes may explain the difference between central and marginal *T*_PM_, because the marginal pit membrane area could be more compressed by deflection against the pit border than the central area that facing the aperture. Nevertheless, this difference raises questions about the assumption that cellulose fibres are homogeneously and equally spaced from each other. It seems likely that the slightly negatively charged cellulose fibres repel each other and are more loosely arranged in the centre (Zhang *et al.*, 2016), but are more compressed near the edge, where the cellulose fibres are firmly anchored into the pectin-rich annulus and primary wall. Although the orientation of microfibrils may not be completely random and appears to be directed by a dual guidance mechanism (Chan & Coen, 2020), it seems unlikely from a developmental point of view that more cellulose fibrils are deposited in the centre than near the annulus, as could be shown for torus-bearing angiosperms (Dute, 2015)

### How is the size of pore constrictions linked to embolism spreading and resistance?

Embolism spreading via pit membranes seems to depend strongly on *T*_PM_, which controls the narrowest pore constriction within a pore. Pit membranes are not different from other non-woven, fibrous porous media, where the pressure required to force a gas bubble through the medium, the so-called bubble point, is a function of the thickness of the medium and its overall structure (Aydilek *et al.*, 2007). Comparison of modelled air-seeding pressures with measurements of *P*_12_ show strong agreement, but a clear difference in absolute values for most species (Fig. 5), with *P*_12_ values falling between the estimated air-seeding based on *R*_*MIN_mean*_ and *R*_*MIN_max*_ (Fig. 4a, 5b, d). Experimental data on air-seeding pressure of angiosperm xylem suggest values between 0.4 and 2 MPa (Choat *et al.*, 2004; Jansen *et al.*, 2009; Christman *et al.*, 2012; Wason *et al.*, 2018), which is more or less in line with *P*_12_ values of a wide range of angiosperm species (Bartlett *et al.*, 2016). Moreover, 65% of the species in our study show *P*_12_ values that are more negative than −2 MPa, with an average *P*_12_ value of −2.57 MPa, which matches the average *P*_12_ value of −2.65 MPa of 12 temperate angiosperm species (Schuldt *et al.*, 2020).

Embolism propagation across thin pit membranes seems to be determined by pores similar in size to *R*_MIN_max_ due to their large similarity between measurements of *P*_12_ and *P*_50_ with modelled air-seeding pressures based on *R*_MIN_max_. In contrast, embolism spreading in species with thick pit membranes is affected by pore sizes that can be close to both *R*_MIN_max_ and *R*_MIN_mean_ (Fig. 3b, 4a). This finding is in line with the fact that high values of *T*_PM_mean_ show a higher standard deviation than low *T*_PM_mean_, while the slope of vulnerability curves becomes lower for species with thicker pit membranes. In fact, *R*_MIN_mean_ is expected to provide an upper limit for air-seeding pressure, since it is unlikely that pore constrictions smaller than *R*_MIN_mean_ will influence air-seeding. Accordingly, *R*_MIN_max_ offers the least resistance to gas moving through a pore space, and provides a good explanation for a lower limit for air-seeding pressure.

There can be various reasons why modelled air-seeding pressures do not match the absolute values of measured *P*_12_ values. First, the values obtained from Model 1 are based on air-seeding estimations of a single pit membrane model with a certain thicknesses, while *P*_12_- and *P*_50_-values represent hydraulically-weighted losses of conductivity at the the vessel network level, which is affected by various structural xylem parameters, such as vessel grouping and the ratio of *T*_PM_ and pit membrane area (Levionnois *et al.*, in press). Second, estimations based on the Young-Laplace equation should be interpreted with caution due to various poorly known parameters and processes. Embolism formation in a multiphase environment under negative pressure is highly complicated, for instance, by dynamic surface tension, line tension, the contact angle of the gas-liquid interface within the pit membrane, and highly variable pore sizes (Choat *et al.*, 2004; Law *et al.*, 2017; Schenk *et al.*, 2017; Satarifard *et al.*, 2018; Zhang *et al.*, 2020; Li *et al.*, 2020; Yang *et al.*, 2020). Pore constrictions and porosity could change if pit membranes become deflected and aspirated against the pit border, which could cause pit membrane shrinkage, reduced porosity and constrictivity, or rearrangement of microfibrils (Tixier *et al.*, 2014; Kotowska *et al.*, 2020; Zhang *et al.*, 2017, 2020). Yet, the mechanical properties of pit membranes remain largely unknown (Tixier *et al.*, 2014).

Moreover, it is also possible that drought-induced embolism spreading does not happen via air-seeding, i.e. mass flow of air-water menisci across intervessel pit membranes. The discovery of surfactant-coated nanobubbles in xylem sap could provide an alternative hypothesis, and highlights the importance of amphiphilic, insoluble lipids associated with pit membranes, and bubble snap-off by pore constrictions (Schenk *et al.*, 2015, 2017, 2018, 2020; Kaack *et al.*, 2019; Park *et al.*, 2019). Diffusion of gas molecules between an embolised and an adjacent vessel could represent an additional way of gas entry and embolism formation, which might be largely dependent on *R*_MIN_mean_ and less on *R*_MIN_max_ (Guan *et al.*, submitted).

### Pit membrane thickness and the number of intervessel pits have different consequences on embolism resistance

We show that *T*_PM_ is a much stronger determinant of the likelihood of leaky pit membranes than *N*_PIT_ and the total intervessel pit membrane surface area (*A*_P_). Our results do not support the rare pit hypothesis (Wheeler *et al.*, 2005; Sperry *et al.*, 2006) and provide a novel view on the relationship between *N*_PIT_ or *A*_P_ and embolism resistance. Most importantly, our Model 2 shows that *T*_PM_ and *N*_PIT_ affect the likelihood of encountering wide pores differently, with contrasting differences for species with a wide range of *T*_PM_. The effect of *N*_PIT_ on vessel leakiness is limited to a narrow range of critical *T*_PM_-values, depending on the assumptions made in Model 2 (Fig. 6, Fig. S4).

In a general, simplified way, three functional types of pit membranes can be distinguished based on *T*_PM_: (1) a thin, risky type, with relatively large pores, a rather low embolism resistance, and little or no reduced embolism resistance for low values of *N*_PIT_, (2) a thick and very safe pit membrane type, with narrow pores, high embolism resistance, and hardly any reduction of embolism resistance for high *N*_PIT_, and (3) an intermediate pit membrane type, with embolism resistance strongly affected by *N*_PIT_, where *N*_PIT_ or other xylem structural traits could potentially be modified during growth to vary embolism resistance in response to the amount of drought experienced. Unfortunately, exact *T*_PM_-values to define these pit membranes types are unclear. Based on leakiness probabilities that are close to zero for *T*_PM_ > 250 nm (Models 1 and 3), and the decreasing slopes of the measured *P*_50_-values with increasing *T*_PM_, we roughly estimate that *T*_PM_ values of the intermediate type are between 150 and 300 nm. This would correspond to 60% of the species in our data set. Interestingly, the risky and safe pit membranes (types and 2) decouple hydraulic safety from hydraulic connectivity, which is suggested to increase with vessel connectivity of the xylem (Loepfe *et al.*, 2007; Schenk *et al.*, 2008; Espino & Schenk, 2009). Since hydraulic connectivity relates to efficiency, the lack of a trade-off between safety and efficiency at the pit membrane level could be suggested, which provides a novel view on the weak relationship between specific hydraulic conductivity and *P*_50_-values of many angiosperm species (Hacke *et al.*, 2006; Loepfe *et al.*, 2007; Gleason *et al.*, 2016).

Overall, our results indicate that the rare pit hypothesis cannot explain embolism spreading at the whole vessel network since the functional importance of multiple pore constrictions makes it highly unlikely that many vessels contain a leaky pore for a wide range of *T*_PM_. In fact, earlier studies that tested this hypothesis should be considered carefully due to possible artefacts in embolism resistance measurements (Wheeler *et al.*, 2013; Torres-Ruiz *et al.*, 2017). Also, no large pores have ever been found in hydrated pit membranes (Schmid & Machado, 1968; Choat *et al.*, 2003, 2004; Pesacreta *et al.*, 2005; Jansen *et al.*, 2018; Zhang *et al.*, 2020). Finally, plants are unlikely to create failures in the three-dimensional development of their cell walls because the synthesis and deposition of cellulose during primary cell wall development includes highly orchestrated processes by the cytoplasm and its cytoskeleton, which reduces the likelihood of large gaps between cellulose fibrils and/or fibrillar aggregates (Chaffey *et al.*, 1997; Oda & Fukuda, 2013; Bourdon *et al.*, 2017; Sugiyama *et al.*, 2017, 2019).

Further progress in understanding embolism spreading in angiosperm xylem will strongly depend on the development of realistic three-dimensional pit membrane and vessel network models (Gaiselmann *et al.*, 2014; Li et al., 2019), combined with careful simulations of the chemical and physical interactions within a multiphase environment of gas, water, cellulose, and surfactants.

## Acknowledgements

Financial support is acknowledged to SJ by a research grant from the German Research Foundation (JA 2174/5-1; nr. 383393940), to SJ and VS by the Baden-Württemberg Ministerium für Wissenschaft, Forschung und Kunst (project 7533-7-11.10-16), and to HJS and SJ by the National Science Foundation (IOS-1754850). We thank Klaus Körber from the Bavarian State Institute for Viticulture and Horticulture, Veitshochheim, Germany, for granting us access to the Stutel-Arberetum facility, as well as Andreas Lösch and all others involved in the ‘Klimabäume Stutel’ project. We thank various colleagues for fruitful discussions, valuable suggestions, and practical assistance in the lab.

## Author contributions

LK, MW, LP, HJS, VS, SJ planned and designed the research. IE, ZK, SL, CT, YZ, BS provided experimental data. LK and MW wrote the manuscript, with input from all co-authors. LK and MW contributed equally.

## Supporting Information

Additional supporting information may be found in the online version of this article.

**Fig. S1** Frequency distribution of the number of intervessel pits per average vessel.

**Fig. S2** TEM images of intervessel pit membranes of different thickness.

**Fig. S3** Results of Model 1, Scenario 2; relation of TPM and pore constriction size.

**Fig. S4** Three-dimensional graph based on the risky scenario of Model 2, with 0.5 probability of having a large pore in a single pit membrane layer.

**Fig. S5** Two-dimensional graph based on Model 2 showing the probability of a large pore in a vessel of up 400,000 pits per vessl.

**Table S1** Dataset of the 31 angiosperm species studied, with reference to the anatomical and hydraulic traits measured.

**Methods** S1 R script of Model 3

**Methods S2** Protocols: plant material, xylem embolism resistance, transmission electron microscopy, vessel and pit dimension

## Notes

### Competing Interest Statement

The authors have declared no competing interest.

