## Supplementary material for "Pore constrictions in intervessel pit membranes reduce the risk of embolism spreading in angiosperm xylem": Fig. S1

The following Supporting Information is available for this article:

**Fig. S1** Frequency distribution of the number of intervessel pits per average vessel.

**Fig. S2** TEM images of intervessel pit membranes of different thickness.

**Fig. S3** Results of Model 1, Scenario 2; relation of $T_{PM}$ and pore constriction size.

**Fig. S4** Three-dimensional graph based on the risky scenario of Model 2, with 0.5 probability of having a large pore in a single pit membrane layer.

**Fig. S5** Two-dimensional graph based on Model 2 showing the probability of a large pore in a vessel of up 400,000 pits per vessel.

**Table S1** Dataset of the 31 angiosperm species studied, with reference to the anatomical and hydraulic traits measured.

**Methods S1** R script of Model 3

**Methods S2** Protocols: plant material, xylem embolism resistance, transmission electron microscopy, vessel and pit dimensions

**Fig. S1** Frequency distribution of the number of intervessel pits per average vessel for 72 angiosperm tree species of 16 families, which varied asymmetrically from 510 to 370,755, and was calculated by dividing the total intervessel pit membrane area per vessel by the average area of intervessel pit membranes. Data are based on multiple data sets (Wheeler et al., 2005; Jansen *et al.*, 2011; Lens *et al.*, 2011; Nardini *et al.*, 2012; Scholz *et al.*, 2013; Klepsch *et al.*, 2016; and original data).

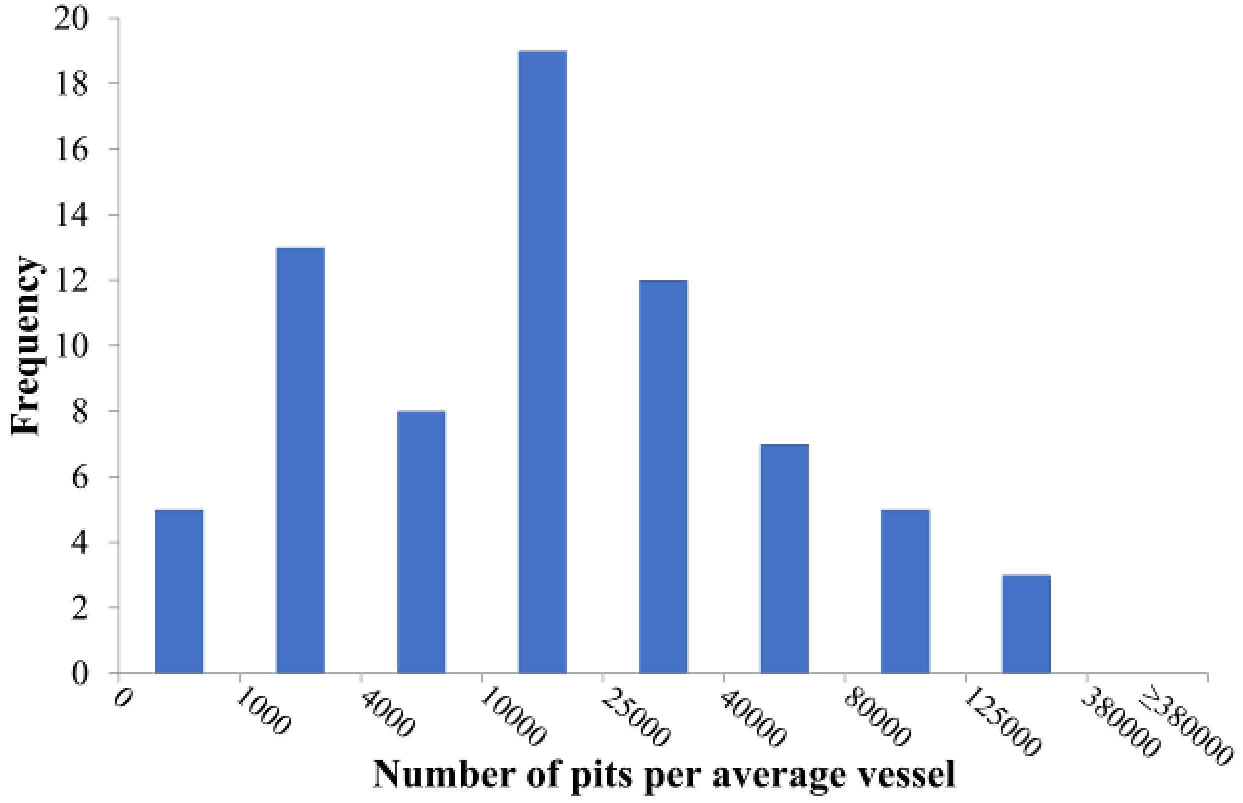

**Fig. S2** Transmission electron microscope images of transverse sections of intervessel pit membranes in woody branches of *Laurus nobilis* (a) and *Acer pseudoplatanus* (b) after fixation with glutaraldehyde and post-fixation with $OsO_4$. Pit membranes of *L. nobilis* are much thicker than *A. pseudoplatanus*, with mean thickness values of 552 nm (± 113 SD) and 270 nm (± 44 SD), respectively. PA = pit aperture; PB = pit border; PC = pit chamber; PM = pit membrane. The white triangles point to the pit membrane thickness, and white arrows show the pit membrane annulus.

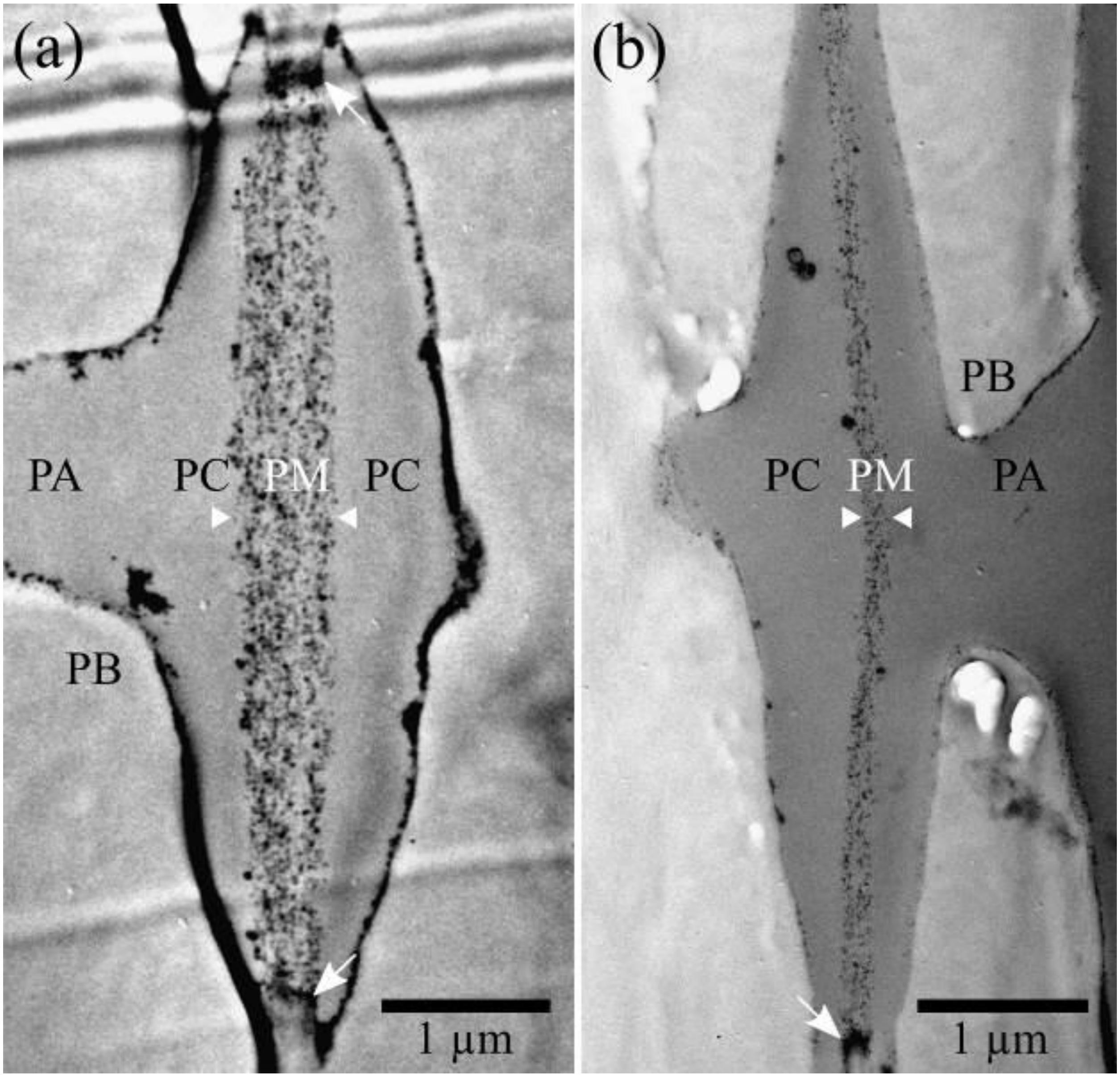

**Fig. S3** Results of Model 1, showing the pit membrane thickness plotted versus the pore constriction diameter based on Model 1. A random number model was used, with the mean pore constriction size set to 100 ± 80 nm, and a minimum size of 5 nm. Pore constriction sizes were determined for 1,100 simulated pores corresponding to an average sized pit membrane.

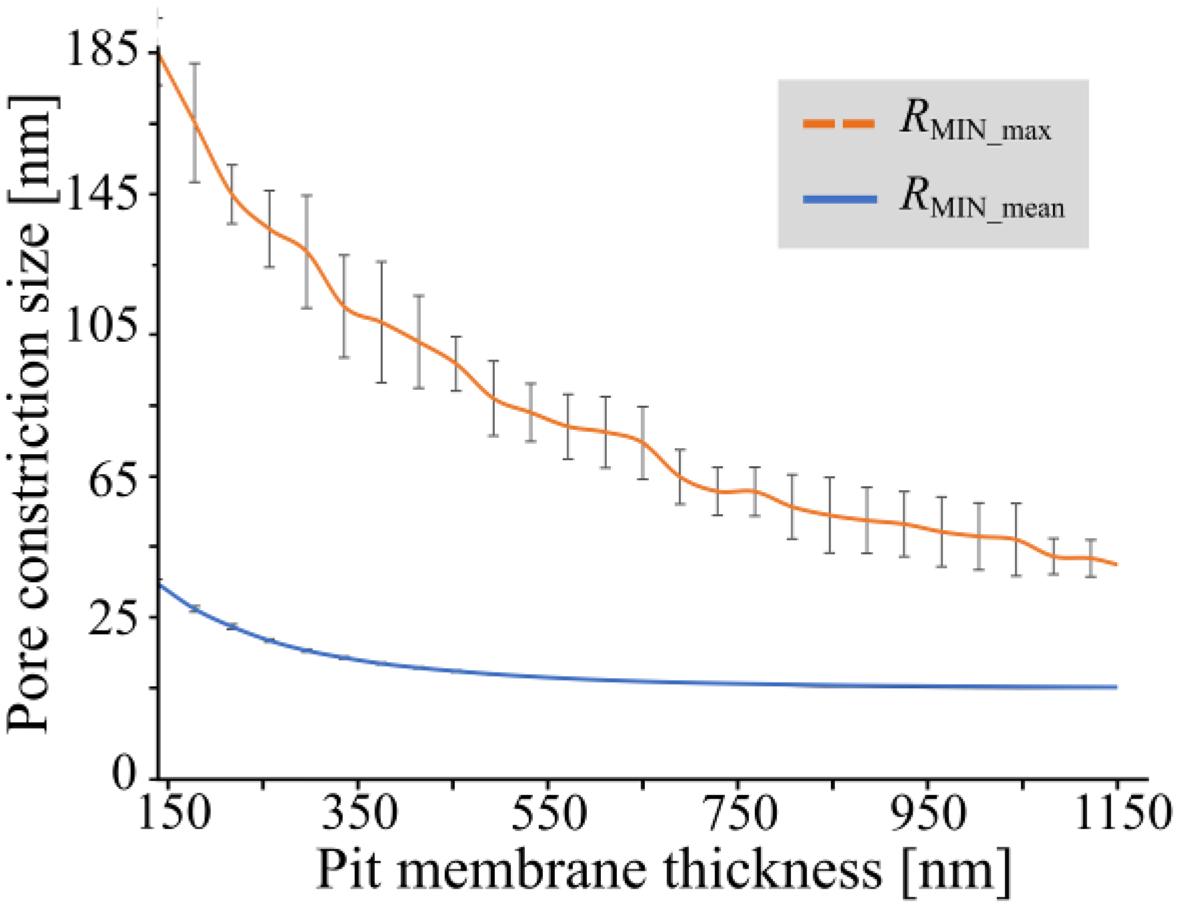

**Fig. S4** The probability of encountering large pores in intervessel pit membranes for an entire vessel decreases with increasing pit membrane thickness as predicted by Model 2. The pit membrane thickness varied from 140 nm to 1,180 nm, and the number of pits per intervessel wall varied between zero to 81,000. The chance of having a large hole within a single microfibril layer was assumed to be 0.5. This model did not consider the actual size of the pore constriction, and ignored whether or not a hole was aligned with other holes in adjacent membrane layers. The green area indicates where most angiosperm species occur based on the number of intervessel pits per vessel, with the median (red dotted line), and the first and third quartile (yellow dotted line).

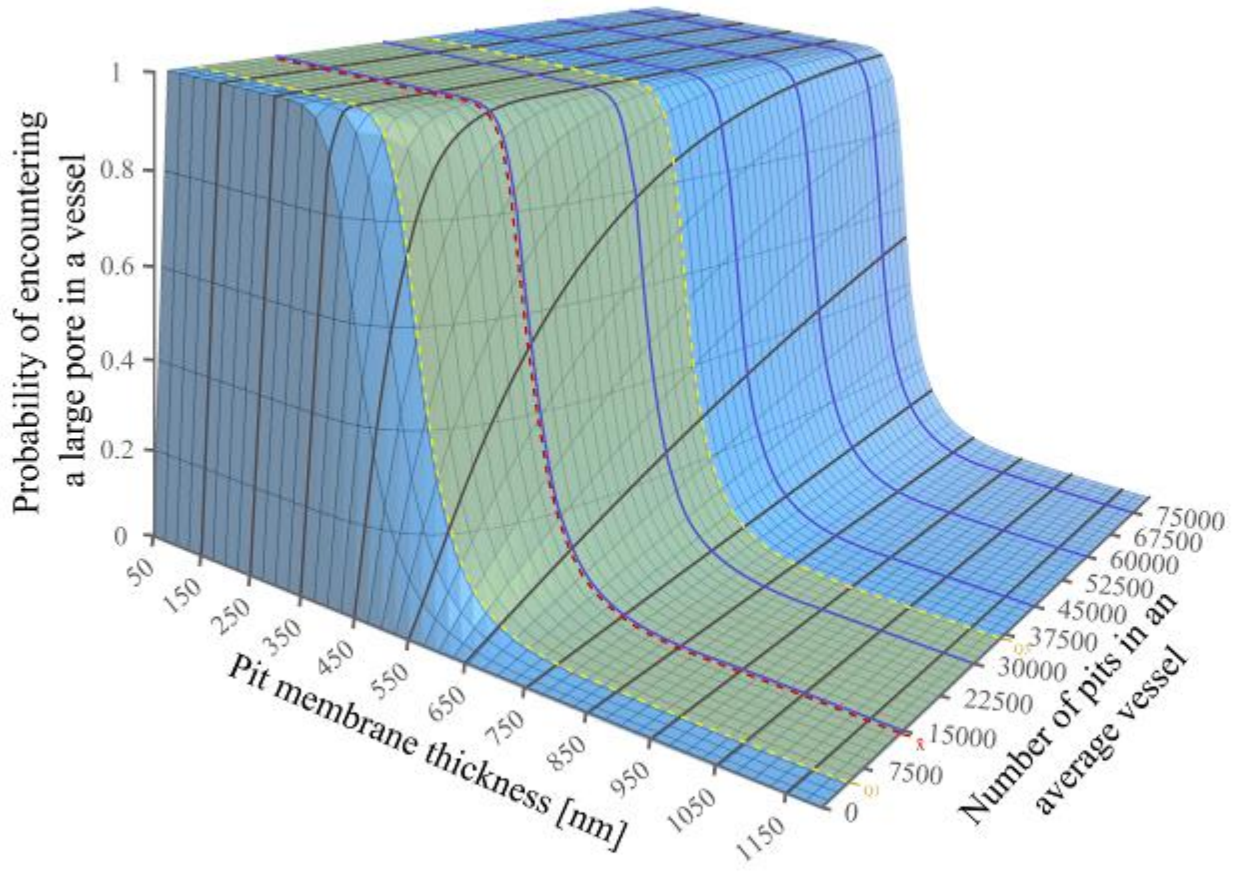


**Fig. S5** The probability of encountering large pores in intervessel pit membranes for a given pit membrane thicknesses over a range of 0 to 400,000 pits per vessel, as predicted by Model 2. The chance of having a large hole within a single microfibril layer was assumed to be 0.25 (a) or 0.5 (b).

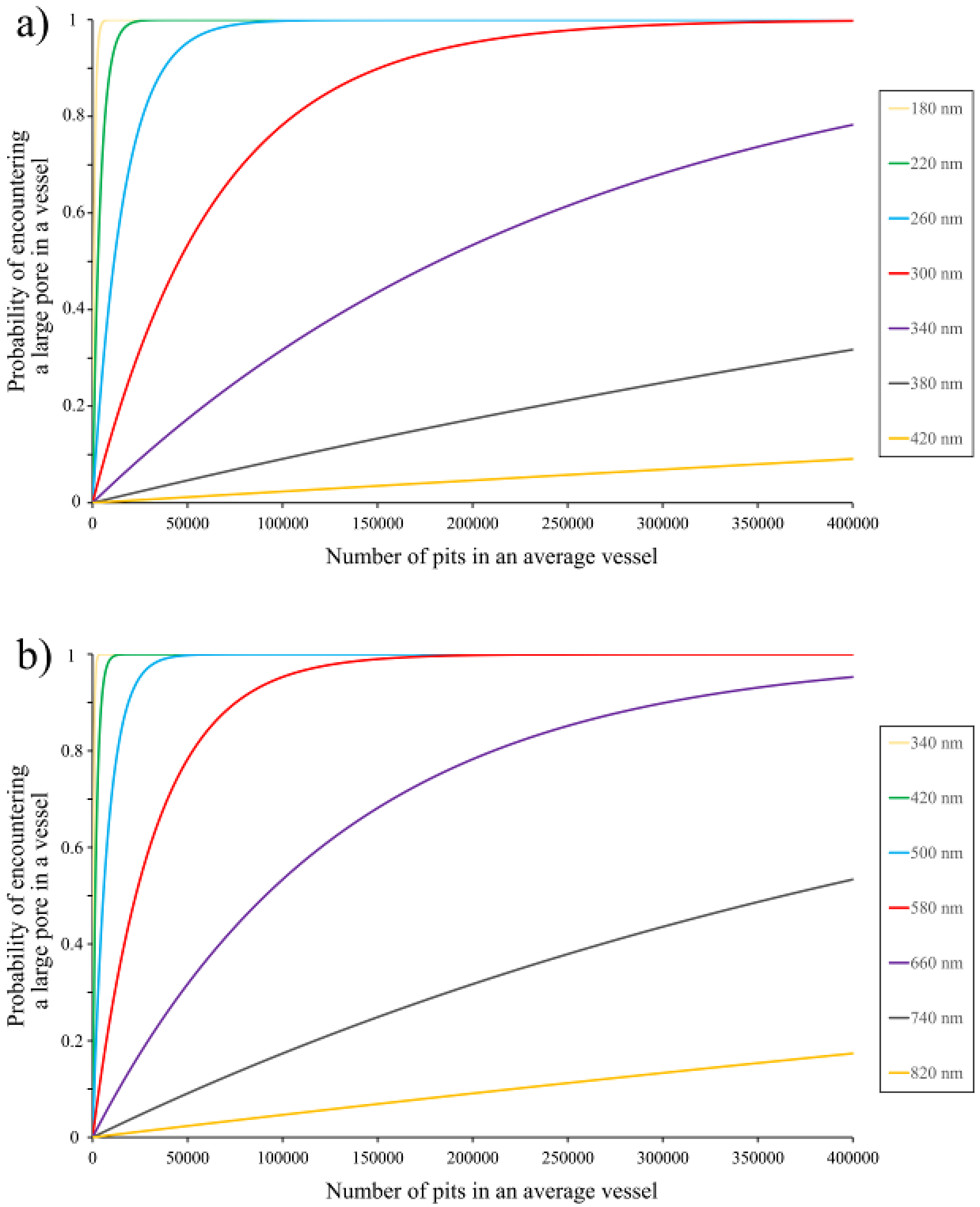
a)
Probability of encountering a large pore in a vessel
Number of pits in an average vessel
0
0.2
0.4
0.6
0.8
1
0
50000
100000
150000
200000
250000
300000
350000
400000
180 nm
220 nm
260 nm
300 nm
340 nm
380 nm
420 nm
b)
Probability of encountering a large pore in a vessel
Number of pits in an average vessel
340 nm
420 nm
500 nm
580 nm
660 nm
740 nm
820 nm

**Table S1** List of the 31 angiosperm species studied, with reference to their family classification, the xylem water potential that corresponds to 12%, 50% and 88% loss of maximum hydraulic conductivity (P12/50/88, MPa), slope of the vulnerability curve, the P50 method, total intervessel pit membrane area per vessel (AP), the thickness of intervessel pit membranes (TPM, nm), and the number of intervessel pit membranes measured on TEM images (N). P50/12/88 of the following species are based on literature: Laurus nobilis (Jansen et al., 2011; Lamarque et al., 2018), Quercus ilex (Jansen et al., 2011; Lobo et al., 2018), Olea europaea (Jansen et al., 2011; Torres-Ruiz et al., 2017), Prunus avium (Cochard et al., 2008; Scholz et al., 2013), and Quercus robur (Choat et al., 2016). All AP values represent original data, except for Laurus nobilis, Quercus ilex, and Olea europaea (Jansen et al., 2011), and Prunus avium (Scholz et al., 2013). Species are arranged from most negative to least negative P50 values. 1 = microCT, 2 = cavimille, 3 = cavitron, 4 = ChinaTron.

| Family | Species | $P_{50}$ [MPa] | $P_{50}$ SE [MPa] | $P_{12}$ [MPa] | $P_{12}$ SE [MPa] | $P_{88}$ [MPa] | $P_{88}$ SE [MPa] | Slope | Slope SE | $A_P$ [mm$^2$] | $T_{PM}$ mean [nm] | $T_{PM}$ mean SD [nm] | $T_{PM}$ centre [nm] | $T_{PM}$ centre SD [nm] | $T_{PM}$ edge [nm] | $T_{PM}$ edge SD [nm] | N | $P_{50/12/88}$ reference, method | $A_P$ reference |
|---|---|---|---|---|---|---|---|---|---|---|---|---|---|---|---|---|---|---|---|
| Lauraceae | *Laurus nobilis* | -7.94 | 0.47 | -5.37 | 0.32 | -10.51 | 0.62 | 19.48 | 6.13 | 0.746 | 552 | 113 | 675 | 151 | 428 | 91 | 18 | Lamarque et al.(2018), 1 | Jansen et al. (2011) |
| Fagaceae | *Quercus ilex* | -7.13 | 0.16 | -4.93 | 0.29 | -9.33 | 0.23 | 24.98 | 2.24 | 0.083 | 476 | 117 | 609 | 170 | 343 | 84 | 19 | Lobo et al. (2018), 2 | Jansen et al. (2011) |
| Rosaceae | *Crataegus persimilis* | -5.91 | 0.17 | -4.66 | 0.49 | -7.17 | 0.14 | 46.21 | 13.07 | | 335 | 70 | 417 | 112 | 254 | 51 | 16 | 3 | |
| Oleaceae | *Olea europaea* | -5.70 | | -3.44 | | -6.27 | | 39.18 | | 0.349 | 610 | 79 | 758 | 104 | 461 | 74 | 35 | Torres-Ruiz et al. (2017), 1 | Jansen et al. (2011) |
| Rosaceae | *Sorbus latifolia* | -5.68 | 0.37 | -3.73 | 0.72 | -7.63 | 0.06 | 27.58 | 5.47 | | 285 | 49 | 325 | 60 | 246 | 44 | 16 | 3 | |
| Sapindaceae | *Acer monspessulanum* | -5.62 | 0.32 | -4.93 | 0.29 | -6.32 | 0.47 | 54.29 | | 0.296 | 348 | 64 | 459 | 111 | 236 | 39 | 16 | 3 | original data |
| Rosaceae | *Pyrus calleryana* | -5.55 | 0.15 | -3.98 | 0.51 | -7.12 | 0.25 | 35.05 | 7.05 | | 303 | 74 | 372 | 96 | 234 | 58 | 9 | 3 | |
| Sapindaceae | *Acer campestre* | -4.76 | 0.67 | -4.49 | | -5.83 | | 35.51 | n.a. | 0.221 | 313 | 25 | 364 | 40 | 263 | 19 | 27 | 4 | original data |
| Rosaceae | *Prunus avium* | -4.76 | | -3.88 | | -5.65 | | 42.70 | 3.15 | 0.152 | 437 | 46 | 525 | 68 | 349 | 30 | 7 | Cochard et al. (2008), 3 | Scholz et al. (2013) |
| Betulaceae | *Ostrya carpinifolia* | -4.56 | 0.17 | -3.83 | 0.20 | -5.29 | 0.17 | 69.53 | 6.11 | 1.450 | 319 | 47 | 360 | 55 | 278 | 48 | 10 | 3 | original data |
| Sapindaceae | *Acer platanoides* | -4.23 | 0.09 | -3.20 | 0.21 | -5.25 | 0.23 | 37.25 | | 0.439 | 327 | 32 | 405 | 42 | 250 | 27 | 15 | 3 | original data |
| Betulaceae | *Ostrya virginiana* | -4.21 | 0.12 | -2.56 | 0.27 | -5.87 | 0.14 | 32.41 | 5.14 | 1.199 | 309 | 29 | 356 | 43 | 261 | 26 | 15 | 3 | original data |
| Fagaceae | *Quercus robur* | -4.16 | 0.16 | -2.38 | | -5.60 | | 26.39 | | | 271 | 60 | 338 | 94 | 204 | 35 | 18 | Choat et al. (2016), 1 | |
| Rosaceae | *Prunus serrulata* | -4.01 | 0.23 | -2.84 | 0.34 | -5.19 | 0.16 | 43.63 | 4.77 | | 264 | 51 | 306 | 71 | 221 | 36 | 12 | 3 | |

| | | | | | | | | | | | | | | | | | | | |
|---|---|---|---|---|---|---|---|---|---|---|---|---|---|---|---|---|---|---|---|
| Betulaceae | *Carpinus betulus* | -3.74 | 0.07 | -2.95 | 0.16 | -4.52 | 0.04 | 66.79 | 6.67 | 0.925 | 315 | 37 | 374 | 58 | 255 | 27 | 16 | 3 | original data |
| Betulaceae | *Ostrya japonica* | -3.49 | 0.05 | -2.65 | 0.10 | -4.33 | 0.08 | 61.82 | 6.02 | 1.237 | 250 | 33 | 282 | 47 | 217 | 30 | 38 | 3 | original data |
| Malvaceae | *Tilia platyphyllos* | -3.39 | 0.06 | -2.81 | 0.18 | -3.98 | 0.09 | 95.55 | 23.35 | | 165 | 18 | 166 | 25 | 164 | 19 | 13 | 3 | |
| Malvaceae | *Tilia tomentosa* | -3.29 | 0.01 | -2.63 | 0.07 | -3.94 | 0.05 | 77.65 | 7.09 | | 218 | 15 | 266 | 25 | 169 | 10 | 7 | 3 | |
| Malvaceae | *Tilia cordata* | -3.11 | 0.15 | -2.25 | 0.22 | -3.97 | 0.14 | 60.46 | 8.76 | | 213 | 24 | 233 | 33 | 193 | 20 | 6 | 3 | |
| Betulaceae | *Carpinus japonica* | -2.94 | 0.04 | -2.24 | 0.01 | -3.63 | 0.07 | 72.72 | 3.41 | 0.730 | 211 | 35 | 241 | 43 | 181 | 31 | 17 | 3 | original data |
| Malvaceae | *Tilia mongolica* | -2.92 | 0.03 | -2.15 | 0.04 | -3.70 | 0.08 | 64.80 | 4.19 | | 222 | 37 | 257 | 44 | 187 | 32 | 13 | 3 | |
| Sapindaceae | *Acer pseudoplatanus* | -2.74 | 0.54 | -2.01 | | -3.53 | | 48.10 | | 0.920 | 270 | 44 | 326 | 64 | 214 | 28 | 14 | 4 | original data |
| Betulaceae | *Betula utilis* | -2.06 | 0.09 | -1.85 | 0.07 | -2.28 | 0.12 | 244.35 | 38.24 | 0.660 | 239 | 65 | 262 | 68 | 216 | 69 | 3 | 3 | original data |
| Betulaceae | *Corylus avellana* | -2.02 | 0.11 | -1.27 | 0.18 | -2.77 | 0.38 | 112.83 | 32.32 | 0.843 | 395 | 61 | 453 | 82 | 395 | 61 | 15 | 3 | original data |
| Platanaceae | *Platanus orientalis* | -1.83 | 0.06 | -1.33 | 0.17 | -2.34 | 0.06 | 111.82 | 29.98 | | 252 | 29 | 313 | 32 | 191 | 35 | 10 | 3 | |
| Betulaceae | *Betula pendula* | -1.78 | 0.03 | -1.62 | 0.03 | -1.95 | 0.04 | 310.81 | 19.95 | 1.096 | 205 | 40 | 225 | 50 | 184 | 33 | 27 | 3 | original data |
| Betulaceae | *Betula pubescens* | -1.76 | 0.06 | -1.53 | 0.11 | -1.98 | 0.11 | 301.71 | 68.05 | 1.264 | 202 | 29 | 217 | 38 | 202 | 29 | 32 | 3 | original data |
| Platanaceae | *Platanus acerifolia* | -1.75 | 0.10 | -1.40 | 0.14 | -2.10 | 0.06 | 146.89 | 15.76 | | 293 | 48 | 339 | 68 | 248 | 36 | 10 | 3 | |
| Betulaceae | *Alnus cordata* | -1.72 | 0.03 | -1.48 | 0.04 | -1.97 | 0.02 | 205.27 | 11.42 | 0.360 | 228 | 35 | 254 | 54 | 201 | 29 | 24 | 3 | original data |
| Betulaceae | *Alnus glutinosa* | -1.48 | 0.03 | -1.35 | 0.02 | -1.62 | 0.04 | 371.77 | 24.70 | 0.791 | 170 | 29 | 181 | 34 | 158 | 27 | 18 | 3 | original data |
| Betulaceae | *Alnus incana* | -1.30 | 0.02 | -1.15 | 0.02 | -1.45 | 0.03 | 340.05 | 25.50 | 0.802 | 171 | 27 | 177 | 35 | 165 | 23 | 20 | 3 | original data |

**Methods S1** R script of Model 3

1st Simulate leakiness of a single pit membrane

Simulate a single pit membrane with radius m_radius and n_l layers Each layer comprises n_p randomly located (non-touching) holes of radius p_radius. This function returns 1, if a sequence of properly aligned pores exists within the simulated pit membrane and 0 otherwise.

a) Parameters

m_radius pit membrane radius
p_radius radius of pore/hole
n_p number of holes per layer
n_l number of layers per membrane
overlap required overlap between holes in adjacent layers

b) Returns

TRUE, if the simulated membrane is leaky, FALSE otherwise

```
isSinglePitMembranLeaky <- function(m_radius, p_radius, n_p, n_l, overlap) {
  last_locations = matrix()
  for (l in 1:n_l) {
    locations = matrix((runif(2) * 2*m_radius) - m_radius)
    p = 1
    while (p < n_p) {
      xy <- (runif(2) * m_radius) - m_radius/2
      if (min(colSums((locations - xy)^2)) > (2*p_radius)^2) {
        locations = cbind(locations, xy)
        p = p+1
      }
    }
    if (l > 1) {
      new_locations = matrix(nrow=2, ncol=0)
      for (p in 1:n_p) {
        xy <- locations[,p]
        if (min(colSums((last_locations - xy)^2)) <= ((1-overlap)*2*p_radius)^2) {
          new_locations = cbind(new_locations, xy)
        }
      }
      last_locations = new_locations

    } else {
      last_locations = locations
    }

    if (ncol(last_locations) == 0) {
      return(FALSE)
    }
  }
  return(TRUE)
}
```

2nd Simulate ratio of leaky pit membranes

Simulate the probability that a pit membrane is leaky, using isSinglePitMembraneLeaky.

a) Parameters

n number of simulation runs
m_radius pit membrane radius
p_radius radius of pore/hole
n_p number of holes per layer
n_l number of layers per membrane
overlap required overlap between holes in adjacent layers

b) Returns

Proportion of leaky pit membranes.

```
proportionOfLeakyPitMembranes <- function(n, m_radius, p_radius, n_p, n_l, overlap) {
  leakyPMs <- 0

  for (i in 1:n) {
    if(isSinglePitMembranLeaky(m_radius, p_radius, n_p, n_l, overlap)) {
      leakyPMs = leakyPMs + 1
    }
  }

  return(leakyPMs/n)
}
```

3rd Example

```
# Parameters
m_radius <- 2500    # Pit membrane radius in nm
p_radius <- 100     # Large pore radius in nm
n_p <- 5            # Number of large pores per layer of pit membrane
n_ls <- c(3, 4, 5, 6, 7, 8, 9, 10, 11, 12)  # Number of layers per pit membrane

n <- 1000       # Number of simulations

overlap <- 0 # How much do holes in adjacent layers need to overlap to be connected? 0 = touching, 1 = completely overlapping

proportionOfLeakyPitMembranes(n, m_radius, p_radius, n_p, n_ls[1], overlap)

## [1] 0.018
```

The mean number of leaky pit membranes, containing at least one leaky pore, can be estimated by multiplying the proportion of leaky pit membranes with the average number of pits in a vessel.

**Methods S2** Protocols: Plant material, Xylem embolism resistance, Transmission electron microscopy, Vessel and pit dimensions

**Plant material studied**

Stem wood samples of 31 species were studied (Table S1), including a large phylogenetic sampling of mainly temperate species. Selection criteria were the availability of plant material to obtain $T_{PM}$ measurements based on fresh samples, and xylem embolism resistance data from vulnerability curves. Samples included healthy branches, which were two to five years old, and few branches that were up to ten years. Tension wood was avoided as much as possible. We also excluded species with a maximum vessel length longer than the diameter of the centrifuge rotor used (i.e. ± 27 cm), since these species would be problematic for the flow-centrifuge method (Wang *et al.*, 2014; Torres-Ruiz *et al.*, 2017; Peng *et al.*, 2019).

Material from 12 species was sampled between June and August 2019 at the Versuchsbetrieb für Obstbau und Gartengehölze of the Landesanstalt für Weinbau und Gartenbau at Veitshöchheim (Germany), and samples of *Acer platanoides* and *A. monspessulanum* were collected at the University of Würzburg. We also included branch material from ten Betulaceae species, which were retrieved from Li *et al.* (2016), and two additional species of *Acer* (*A. pseudoplatanus*, *A. campestre*). The Betulaceae species were collected at Ulm University in May 2013, and the two *Acer* species in July 2019. All samples from Veitshöchheim and Würzburg included paired samples, with $T_{PM}$ and embolism resistance measurements based on the same branches. For samples collected at Ulm, embolism resistance and anatomy were based on different stem samples from the same shrub or tree.

To cover a wider range of embolism resistance, we added five species from literature that were highly embolism resistant: *Laurus nobilis*, *Olea europaea*, *Prunus avium*, *Quercus robur*, and *Q. ilex*. Material of these five embolism resistant species could not be paired, and was based on samples from different specimens. Plant material for $T_{PM}$ measurements of these species came from the botanical garden of Ulm University. Interspecific variation in $T_{PM}$ and embolism resistance, however, is generally larger than intraspecific variation (Jansen *et al.*, 2009; González-Muñoz *et al.*, 2018; Kotowska *et al.*, 2020). Moreover, seasonal variation between the samples collected was ignored.

### Xylem embolism resistance

Xylem embolism resistance was measured by estimating the xylem water potential (MPa) that corresponded to 50% loss of maximum hydraulic conductivity, i.e. the $P_{50}$ value (MPa). We used a Cavitron (Cochard *et al.*, 2005) for a total of 24 species, and a ChinaTron centrifuge (XiangYi Centrifuge Instrument Co., Ltd., Changsha, China) to obtain $P_{50}$ values for *A. campestre* and *A. pseudoplatanus*. The methodology of both flow centrifuges was similar. Moreover, $P_{12/50/88}$ values and the slope of vulnerability curves were taken from literature based on microCT observations of *Q. robur* (Choat *et al.*, 2016), *L. nobilis* (Lamarque *et al.*, 2018), and *O. europaea* (Torres-Ruiz *et al.*, 2017), and the flow-centrifuge technique for *P. avium* (Cochard *et al.*, 2008), and *Q. ilex* (Lobo *et al.*, 2018). Comparison of microCT based vulnerability curves the with flow-centrifuge approach showed that both methods can be compared when the open-vessel artefact is avoided (Choat *et al.*, 2016; Torres-Ruiz *et al.*, 2017; Lamarque *et al.*, 2018).

For each species, five branches with a similar basal diameter (ca. 8 mm) were transported to the lab at Göttingen University in 2013, or Würzburg University and Ulm University in 2019. Demineralised, filtered (0.2 µm) and degassed water with 10 mM KCl and 1 mM $CaCl_2$ was used for all hydraulic measurements. Branches were stored in a refrigerated dark room at 4°C, and all the measurements were performed within one week after sample collection. The stem segments were recut under water to 27.8 cm, and the bark was removed up to ca. 5 cm from both stem ends and segments inserted into a custom-made rotor chamber. At Würzburg University, long stem samples were recut under water in order to relax the xylem pressure before centrifuge measurements. Measurements started at a negative pressure of -0.834 MPa, which was stepwise lowered until the percentage of loss of conductivity (PLC) reached at least 90%. Vulnerability curves were generated by plotting PLC against xylem pressure, and the $P_{50}$-value was calculated according to a sigmoidal function (Pammenter & Van der Willigen, 1998), and a Weibull function was applied to the Betulaceae species from Ulm. Subsequently, the xylem pressures causing 12% and 88% loss of conductivity ($P_{12}$ and $P_{88}$) were calculated following Domec and Gartner (2001).

### Transmission electron microscopy

All TEM samples were prepared at Ulm University, except for the 12 species collected at Veitshöchheim, which were prepared at Würzburg University. A slightly different TEM preparation procedure was applied between the samples embedded at both universities. Small xylem slivers were cut with a fresh razor blade from the two outmost growth rings of fresh wood samples. These were recut into small cubes (1x2x2 mm) in water. The samples prepared at Ulm were washed several times with PBS (Phosphate-Buffer Saline solution) for five to ten minutes. Specimens at Ulm were then fixed with a standard solution (2.5% glutaraldehyde, 0.1 mol phosphate, 1% sucrose, pH 7.3), post-fixed with 2% aqueous $OsO_4$ for one to two hours at room temperature, stained *en bloc* with uranyl acetate, and dehydrated through a gradual ethanol series (30%, 50%, 70%, and 90%) for two to three minutes. Samples at Ulm were embedded in Epon resin (Sigma-Aldrich, Steinheim, Germany) at 60°C.

Samples at Würzburg were fixed with 5% glutaraldehyde in a 0.1M cacodylate buffer solution, washed several times in 50 mM cacodylate buffer for five minutes, and post-fixed with a 2% $OsO_4$ solution in ice. The samples were then washed five times in distilled water for three minutes each, placed in uranylacetate overnight at room temperature, dehydrated in a gradual ethanol series, treated with 100% propylenoxid, and gradually embedded in Spurr resin, which was polymerized at 60°C and under a partial vacuum of 58 kPa for 48 hours. Although it is unlikely that these different preparation protocols had an effect on $T_{PM}$ as seen under TEM, minor differences cannot be fully excluded (Kotowska *et al.*, 2020).

Ultra-thin sections with a thickness of 60 nm and 100 nm were cut with a diamond knife using an ultra-microtome (Leica Ultracut UCT, Leica Microsystems, Vienna, Austria) and deposited on a copper grid (Athena, Plano GmbH, Wetzlar, Germany). Observations were carried out using a JEM-1210 TEM (Jeol, Tokyo, Japan) at an accelerating voltage of 120 kV at Ulm, and 200 kV at Würzburg. Digital images were taken using a MegaView III camera (Soft Imaging System, Münster, Germany) at Ulm, and a TemCam-F416 camera (TVIPS, Gauting, Germany) at Würzburg.

Image analysis was based on ImageJ (National Institutes of Health, Bethesda, Maryland, USA) to obtain $T_{PM}$ measurements on at least seven intervessel pits per species, although we were able to measure more than 10 pits per species for the majority of species. In various species, intervessel pits were highly transparent, with unclear outlines of the pit membranes. Also, we only included pit membranes when the pit apertures were visible, which indicated that the section was near the centre of the pit membrane. Because pit membranes can be cushion-shaped, $T_{PM}$ was measured separately at the centre and near the edges (i.e. towards the pit membrane annulus). The mean $T_{PM}$ was defined as the mean value of the orthogonal thickness measurements at the centre and near the annulus. Shrunken or aspirated pit membranes, which typically look much darker and thinner under TEM, were excluded (Zhang *et al.*, 2020; Kotowska *et al.*, 2020).

### Vessel and pit dimensions

We estimated the total pit membrane surface area per vessel ($A_P$) and the number of intervessel pits ($N_{PIT}$) per vessel based on anatomical measurements of stem samples for a total of 20 species. These species included 16 of the 31 species that were selected for $T_{PM}$ and $P_{50}$ measurements. Besides 12 Betulaceae species, and four species of *Acer* (Sapindaceae), $A_P$ values were retrieved for four species from literature, namely *Quercus ilex*, *Olea europaea,* and *Laurus nobilis* from Jansen et al. (2011), and Prunus avium from Scholz et al. (2013). $A_P$ estimations from literature followed the same protocol as applied here. Both $A_P$ and $N_{PIT}$ were based on at least three individuals per species. All image analysis measurements related to $A_P$ and $N_{PIT}$ were based on ImageJ. Jansen *et al.* (2011), and *Prunus avium* from Scholz *et al.* (2013)

$A_P$ estimations followed Wheeler *et al.* (2005a) and Hacke *et al.* (2006). Briefly, $A_P$ was determined as the product of the intervessel pit fraction (i.e. the mean fraction of the total vessel wall area occupied by intervessel pits) and the vessel wall area. The intervessel pit fraction was obtained by multiplying the intervessel contact fraction with the ratio of the intervessel wall area occupied by intervessel pits to the total intervessel wall area, which were both measured on light microscopy images of transverse and tangential sections, respectively. The vessel wall area required measurements of the average vessel length and the hydraulically weighted vessel diameter, which was based on a minimum of 300 vessels per species.

Vessel length distribution was based on a minimum of three stems per species using a silicon injection method (Sperry *et al.*, 2005; Wheeler *et al.*, 2005b; Jansen *et al.*, 2011). To avoid embolism, samples of about 28 to 40 cm length were either flushed for 20 minutes using a degassed ultra-filtered solution of distilled water with 10 mM KCl and 1 mM $CaCl_2$, or were immersed in water under vacuum for 48 to 72 hours. A fluorescent silicone mixture (Uvitex and Rhodorsil ESA7250 A + B; Bodo Müller Chemie GmbH, Offenbach, Germany) was injected at the acropetal end of the branch, under a pressure of 0.2 MPa with a pressure chamber (PMS Instrument Company, Albany, OR, USA) for at least four hours. The injected samples were then left on the bench overnight. About five to six transverse sections were cut with a microtome at different distances from the injection point, starting at 6 mm and the final sections were made where only few silicon filled vessels were evident. In each section, the number of vessels filled with silicone was counted using a fluorescent light microscope (Leica DMBRE, Leica Microsystems GmbH, Wetzlar, Germany). Vessel length distributions were determined for complete radial sectors of growth rings to ensure correct representation of intra-ring variation in vessel length. We followed the equation provided in Christman *et al.* (2009) to estimate mean vessel length for each species.

$N_{PIT}$ was estimated by dividing the total intervessel pit membrane area per vessel ($A_P$) by the average intervessel pit membrane area, assuming all pit membranes have a similar diameter and surface area. Species with highly variable intervessel pit sizes, such as the elongated pits in *Vitis vinifera*, were excluded. Pit membrane surface area was calculated based on images of tangential intervessel walls taken with a scanning electron microscope (Zeiss, DSM942, Germany) at an accelerating voltage of 10kV.
